# The network structure and eco-evolutionary dynamics of CRISPR-induced immune diversification

**DOI:** 10.1101/850800

**Authors:** Shai Pilosof, Sergio A. Alcalá-Corona, Tong Wang, Ted Kim, Sergei Maslov, Rachel Whitaker, Mercedes Pascual

**Affiliations:** Department of Life Sciences, Ben-Gurion University of the Negev, Beer-Sheva, Israel; Department of Ecology and Evolution, University of Chicago, 1103 E 57 st, Chicago, 60637, USA; Department of Physics, Loomis Laboratory of Physics, University of Illinois at Urbana-Champaign 1110 West Green St, Urbana, Illinois 61801 USA; Carl R. Woese Institute for Genomic Biology, University of Illinois at Urbana-Champaign, Urbana, Illinois, USA; Department of Microbiology, University of Illinois at Urbana-Champaign, Urbana, Illinois, USA; Department of Bioengineering, University of Illinois at Urbana-Champaign, Urbana, Illinois 61801 USA; Santa Fe Institute, Santa Fe, New Mexico, USA

## Abstract

As a heritable sequence-specific adaptive immune system, CRISPR-Cas is a powerful force shaping strain diversity in host-virus systems. While the diversity of CRISPR alleles has been explored, the associated structure and dynamics of host-virus interactions have not. We explore the role of CRISPR in mediating the interplay between host-virus interaction structure and eco-evolutionary dynamics in a computational model and compare results with three empirical datasets from natural systems. We show that the structure of the networks describing who infects whom and the degree to which strains are immune, are respectively modular (containing groups of hosts and viruses that interact strongly) and weighted-nested (specialist hosts are more susceptible to subsets of viruses that in turn also infect the more generalist hosts with many spacers matching many viruses). The dynamic interplay between these networks influences transitions between dynamical regimes of virus diversification and host control. The three empirical systems exhibit weighted-nested protection networks, a pattern our theory shows is indicative of hosts able to suppress virus diversification. Previously missing from studies of microbial host-pathogen systems, the protection network plays a key role in the coevolutionary dynamics.

## Introduction

Microbial hosts have developed a range of resistance mechanisms against viruses. For example, mutations in receptors leading to surface resistance, innate immunity obtained with restriction-modification systems, and adaptive immunity obtained via Clustered Regularly Interspaced Short Palindromic Repeats (CRISPR) and its CRISPR-associated (Cas) proteins^1–3^. These and other defense mechanisms can operate alone or in tandem to affect the diversification of microbes and viruses^4,5^ as well as to structure the complex infection networks into which this diversity is organized^6–8^. Infection structure (‘who infects whom’) is a crucial feature of microbe-virus interactions because, as for other host-parasite systems, it may affect the evolution^9,10^, population stability^11^ and transmission dynamics^12^ of both partners. Hence, given the crucial role microbes and viruses play in virtually all earth’s ecosystems, infection structure may have far-reaching consequences for ecosystem functions.

Microbe-virus interaction structure is non-random, with two dominant macroscopic topologies in its descriptions: modularity and nestedness^13,14^. Modularity concerns patterns of specificity, in which the network is partitioned into modules of microbes and viruses that interact densely with each other but sparsely with those outside the group. Nestedness reflects instead patterns of specialization in which specialist microbes are infected by subsets of viruses that in turn also infect the more generalist microbes^15^. An analysis of a large assemblage of bacteria-virus infection networks found that these are predominantly nested rather than modular^15^ but a later study suggested that structure depends on phylogenetic scale, with modularity at large phylogenetic scales of species interactions and nestedness among interacting strains of the same species^16^. Several hypotheses have been put forward to explain the emergence of modularity and nestedness in bacteria-phage infection networks from immunity-related coevolutionary dynamics^6,7^ (mutations that create new CRISPR alleles and variation in the relative frequencies of alleles over time). For example, Fortuna et al.^8^ have shown that bacteria-virus networks evolve nestedness under directional arms race dynamics, but not under fluctuating selection.

While providing crucial knowledge about the structure of microbe-virus interactions, all previous studies have focused exclusively on patterns of infection. Infection structure should critically depend however on the genetic basis of resistance of hosts to pathogens^10^. The emergence of network structure from immune selection has been shown in pathogens of humans such as *Plasmodium falciparum*^17,18^. So far, these studies have only analyzed networks of genetic similarity from the parasite perspective thereby inferring the selective impact of hosts because information on host immunity structure and diversity is typically absent. Hence, how immunity structures host-parasite interactions, for example into modular, or nested topologies, is unknown. In contrast, CRISPR is a heritable adaptive immune system conferring sequence-specific protection against viruses, plasmids, and other mobile elements^1,19^. The CRISPR system functions as an adaptive immune system by incorporating DNA segments called ‘protospacers’ of infecting viruses into host genomes as ‘spacers’ that constitute sequence-specific immunity and memory^19^. Hence, it provides a direct sequencebased link that allows consideration of associated structures for both infection and immunity, and from both host and parasite perspectives.

The diversity of CRISPR alleles in natural, experimental, and simulated populations has been explored^20–24^. Simulation studies^20^ followed by experimental tests showed that increased strain diversity of *Pseudomonas aeruginosa* (with strains being clones with different spacer compositions) promotes the ability of the bacterium to control the DMS3vir virus^22^. Nevertheless, how such strain diversity evolves, the emergence of the structure of host-virus interaction networks from molecular mechanisms that link genotype to defensive phenotype, and how networks are assembled and disassembled remain unexplored.

Here, we address how the genetic basis of CRISPR immunity shapes the structure of both infection and protection networks, and in turn, how the structure of the networks shapes the population dynamics of viruses and their hosts. Because we consider not just the presence/absence of protection but the weight or redundancy of the protection link, the two networks are not simply complementary, and the structure of one cannot inform us about that of the other. Using a system of Lotka-Volterra-like differential equations coupled with stochastic evolution (mutation and spacer acquisition), we investigate for the first time the emergence and consequence of the structure of immunity networks. We show that infection and immunity networks have distinct structures emerging from CRISPR immunity that are involved in a constant interplay with the diversification and population dynamics of host and viral strains. This structure influences dynamics with eco-evolution feedbacks.

## Results

### Dynamics shift between virus divesification and host control

Previous work has shown the possibility of diverse populations, containing many strains, with distributed immunity of the host^20,25^. The dynamics of these populations can exhibit two alternating major regimes, escape and diversification of the viruses and dominance of the microbial host, respectively^25^. We extended the model to allow for larger host and virus richness and to track the coevolutionary history of both hosts and viruses through time (Methods). Regime switching is an emergent property of the model and the timing of the switching between the two regimes is defined and identified here based on the dynamics of virus abundance (Methods). In the ‘virus-diversification regime’ (VDR), virus strains proliferate and diversify while in the ‘host-controlled regime’ (HCR) host strains are able to constrain virus diversification and lead to their extinction (Fig. 1). During the former, viruses and hosts coexist with fluctuating abundances, whereas during the latter, hosts reach carrying capacity and viruses exhibit declining abundances and richness followed by either escape or extinction (Figs. S1, S2, S3). Due to the stochastic nature of our simulations, the number of alternations between regimes varies. Nonetheless, for our set of parameters, the long-term dynamics of the system consists of complete virus extinction during one of the HCRs. In most simulations (≈ 70%), viruses do not go rapidly extinct and exhibit instead one or more transient periods of diversification (Fig. 1), whereas in the rest, they show rapid extinction as hosts exert immediate control. Our analyses concentrate on the alternating transient dynamics that precede extinction.

**Fig. 1.**
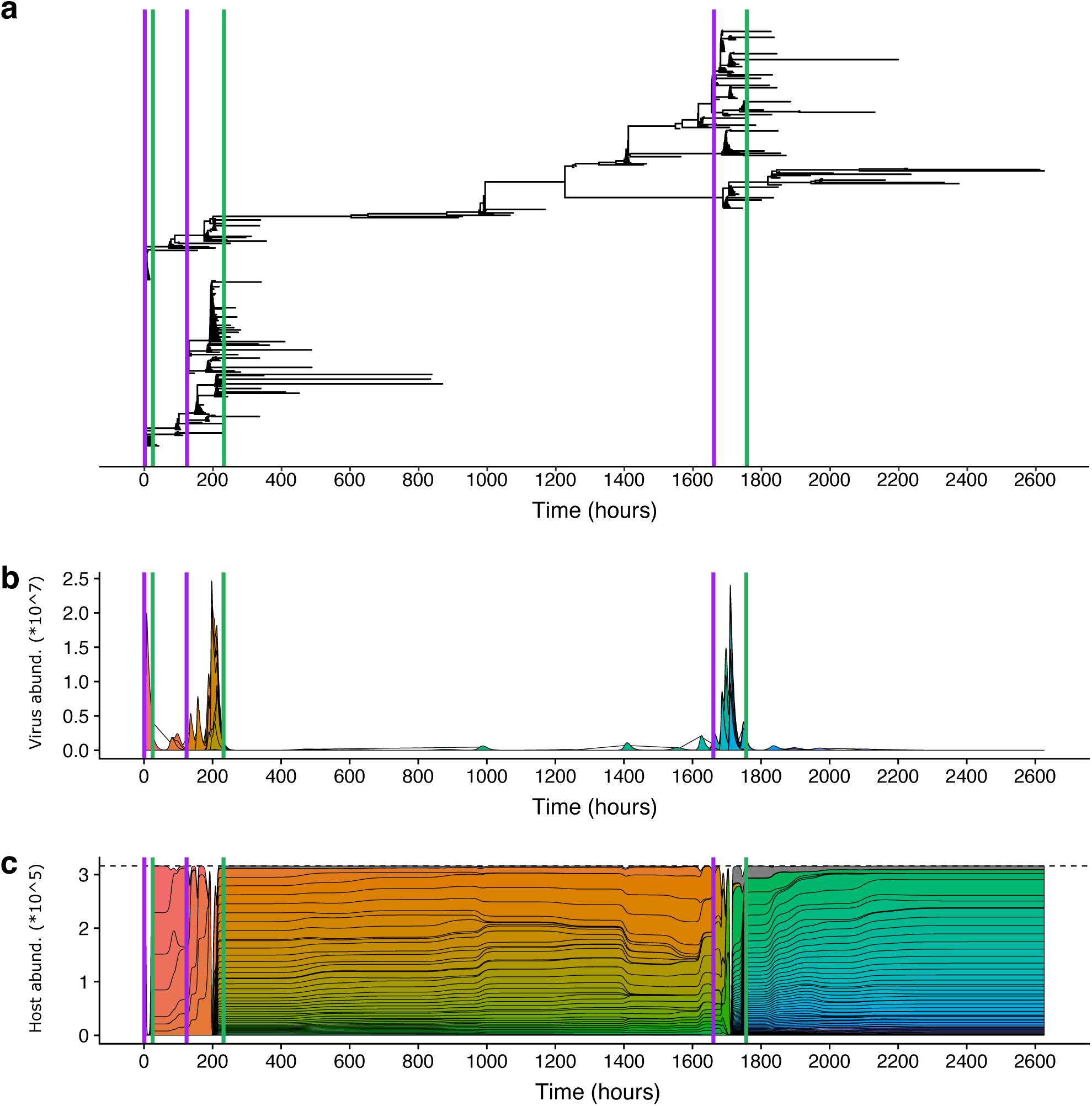
An example of typical dynamics with regime shifts. Virus-diversification regimes (VDRs) start at purple vertical lines and end in the green vertical lines, when the host-controlled regimes (HCRs) begin. **a**, Viral phylogenetic tree shows virus diversification during VDRs and virus extinctions during the HCRs. Intermittent virus diversification in the later part of HCRs eventually leads to escape and the initiation of a new VDR. Complete virus extinction occurs at the end of this simulation during the last HCR. **b** and **c** are the abundance profiles of viruses and hosts, respectively. The 100 most abundant strains are colored, the rest are aggregated and shown in gray. Note that virus growth does not imply a decrease in host richness while host dominance does decrease virus abundance and richness sharply. Virus diversification implies a temporary escape from host control and their rise in abundance, which in turn increases encounters with hosts. The resulting rise in per-capita infection rates of hosts leads to their concomitant diversification because new host strains are generated by acquiring at least one new protospacer. Therefore, despite an initial decline in the abundance of hosts’, their richness typically rebuilds (Fig. S3).

Variation in viral or host abundance alone cannot drive the transitions between these two regimes, as these do not exist in the corresponding Lotka-Volterra dynamics under neutral conditions of hosts not acquiring specific immunity (Supplementary Methods). An alternative explanation is that the structure of strain diversity, emerging from the eco-evolutionary dynamics via specific immunity, explains the transitions between these dynamical regimes. By ‘eco-evolutionary dynamics’ we refer to the dynamic interplay between ecological (e.g., population dynamics such as relative abundances) and evolutionary (e.g., mutation) processes. To investigate the structure of diversity in this complex system, we consider two complementary bipartite networks, the ‘infection’ and ‘immunity’ graphs, through time. In these networks, a node represents a strain of a virus (unique combination of protospacers) or a host (unique combination of spacers), and edges represent a given type of interaction, either infection or protection from infection (Fig. 2).

**Fig. 2.**
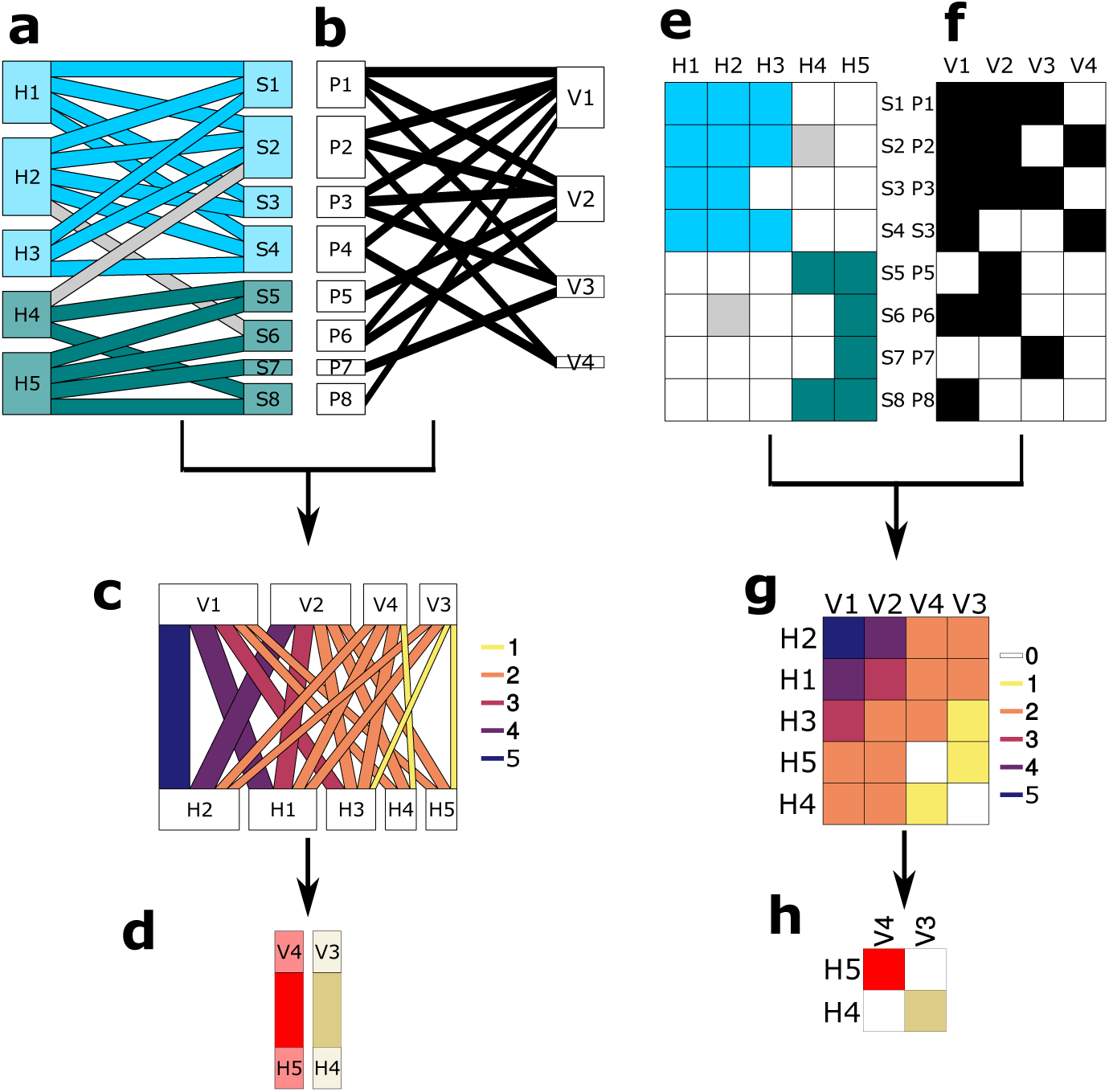
Structures of diversity. Diagrams illustrating the two different kinds of networks (left column) and associated matrices (right column) used in this study and how they are built. The toy example has 5 hosts (H1-H5), 4 viruses (V1-V4), 8 spacers (S1-S8) and 8 protospacers (P1-P2). **a** A bipartite network depicting the spacer composition of hosts. Hosts are affiliated to one of two modules (depicted in light blue and dark green), and interactions can fall either within a module (colored) or outside the module (gray). **b** A bipartite network (not modular) depicting the protospacer composition of viruses. **c** The immunity network is created by counting the number of shared spacers and protospacers between pairs of hosts and viruses (i.e., matches). Interactions in the network are weighted by the number of matches, here depicted by different colors and width. **d** The infection network is created by considering unrealized interactions in the immunity network (equivalent to 0-matches in **g**). Here, only two such interactions exist, between 2 viruses and 2 hosts. There are two modules, depicted in colors, with interactions occurring within the modules only. **e** The counterpart matrix of the network in **a**. Interactions (matrix cells) depict the occurrence of a spacer in a host strain, and are colored as in **a. f** The counterpart matrix of the network in **b**. Matrix cells depict the occurrence of a protospacer in a virus strain. **g** The counterpart matrix of the network in **c**, with colors depicting the number of spacer matches. The matrix is organized by the sum of columns and rows and is quantitatively nested. **h** The counterpart matrix for the network in D. The two interactions correspond to the empty matches in **g**.

### Modularity of infection networks represents niches for hosts

At the start of the simulations and during VDRs, the infection network is built over time by the addition of host and virus strains (Fig. S4). We find that at the end of each VDR, after the infection network has been built, the infection network is significantly modular (*P* < 0.001 in 2 of 3 VDRs; in the first VDR the network was too small to test; Methods). Infections are concentrated within modules of viruses and hosts, with more edges within than between these groups (Fig. 3a). These modules reflect different niches for virus growth whereby hosts are resources. The creation of niches is linked to the structured genetic diversity of the hosts. Specifically, modules in host-spacer networks delineate groups of hosts that share immunity via the same spacers and are therefore susceptible to similar viruses. We find that host-spacer networks aggregated across the VDR are significantly modular (*P* < 0.001 in 2 of 3 VDRs). Aggregation accounts for the fact that hosts accumulate spacers in time without going extinct.

**Fig. 3.**
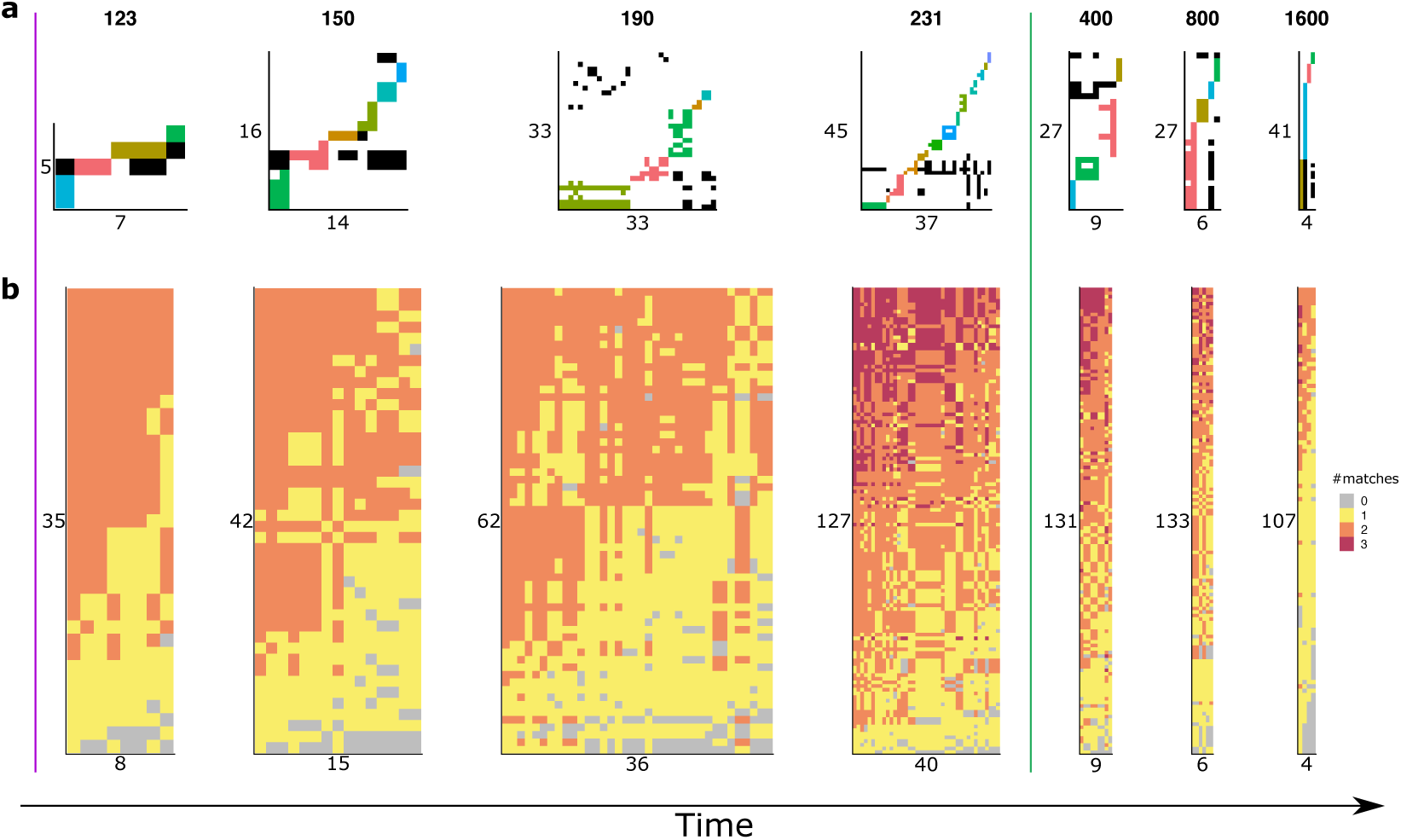
Snapshots of network structure during the two regimes. Networks are defined as in Fig. 2. Each network is a snapshot of the population, with time steps depicted above the networks corresponding to those marked in Fig. 4. Numbers in the x and y axes indicate the number of viruses and hosts participating in each of the networks. Because the size of the matrix is limited, the width and height of the networks are not necessarily proportional to the number of strains (e.g. a reduced number of strains results in panels with larger squares). The purple and green lines depict the initiation of a virus-dominated and host-controlled regimes (VDR and HCR), respectively. **a**, Modularity in the infection network. Colored interactions fall inside modules of virus strains that infect similar hosts (each module has a different color). Black interactions are those that fall outside all modules. The size of the network and the number of modules increase during the VDR (between purple and green vertical lines) and decline during the HCR (right of the green vertical line). **b**, Weighted nestedness in immunity networks. Colors of interactions depict the number of matches between viruses and hosts. The network has a weighted-nested structure, which enforces an orderly viral extinction. Nestedness builds up during the VDR and declines during the HCR, as viruses go extinct. The extinction is orderly, with viruses to which many hosts has immunity via many spacers (those at the left), going extinct first.

The importance of immune selection in the formation of these niches can be demonstrated by asking whether clonal expansion alone could account for the modularity of the infection network. This generally is not the case (i.e., no consistent phylogenetic signal) in the three VDRs both in the infection and host-spacer networks, as shown by comparison of the observed phylogenetic distance within modules to distances obtained in randomized versions of the network (Methods; Tables S1, S2). Moreover, in the absence of host memory, diversification of the viruses and coexistence of different strains is not observed. This is demonstrated by a neutral model in which all the processes of the full system are retained except for the specific memory of the host (Supplementary Methods).

The persistence of the modular structure in the infection network and the coexistence of a diverse community of viral strains is only transitory because the number of susceptible hosts available to all viruses declines rapidly, and no virus is able to infect all hosts. Interactions become organized into the modular structure, with hosts within a module becoming rapidly unavailable for viruses by either lysis or acquisition of protection. This closing of niches coupled with the inability of the viruses to escape makes the VDR effectively transient. To understand why virus escape becomes highly unlikely, we turn next to the structure of the protection network.

### Weighted-nestedness of immunity networks is indicative of host control

Diversification of viruses, enabled by the modular structure of the infection network, increasingly diversifies the host population (Fig. S2). In other words, escape from host control via mutation of protospacers allows higher abundances of particular virus strains, who therefore also experience higher encounter rates with hosts in general. This leads to the increasing acquisition of new spacers through either the failure of their previous immunity or infection by an escape mutant. Such red-queen coevolutionary dynamics (i.e., the reciprocal evolution of hosts and viruses) progressively adds spacers to hosts during the VDR, building the immunity network (Fig. 3b). In this network, edges indicate at least one spacer-protospacer match, and the weight of the edge encodes the number of different matches protecting a given host from a given virus (Fig. 2c). By definition, a lack of an edge indicates infection, and an edge value larger than one indicates redundancy in immunity.

We find that this redundancy has a characteristic quantitative nested structure^26^. We can order the network such that hosts with immunity to most viruses also have more matches to those viruses, and subsequent hosts (from top to bottom) are immune to subsets of viruses, via fewer matches (Fig. 2g, Fig. 3b). Similarly, each virus strain (from left to right) can infect a progressively smaller subset than the virus following it, also via fewer matches. The weighted nested structure is assembled by a complex interplay of temporal changes in abundances, associated encounter rates, and selection pressures. Hosts can increase matches to a virus strain by exposure to other strains that carry an overlapping protospacer, building redundancy. In addition, how the matrix structure is woven from bottom to top, and left to right, with the respective addition of new hosts and viruses, is reflected in the relationship between the order of the rows and columns and strain age (Fig. S5). It is also the result of CRISPR failure, which allows infection and the subsequent acquisition of additional spacers; that is, when a virus infects a protected host whose CRISPR system failed, then the host can (if it does not die) obtain a new spacer. A nested structure can provide universal resistance (when the network is fully connected) and eventually lead to virus extinction. The inability to infect does not mean however immediate or complete extinction of all viruses, because the decline in abundance of virus populations takes time. During this time virus strains can still mutate and therefore escape. How the structure of the protection network implies its own change in ways that ultimately facilitate escape is described next.

### Order in virus extinctions facilitates escape

The nested pattern in the redundancy of immunity, built during a VDR, enforces order and predictability in virus extinctions during the subsequent HCR. Viruses for which most hosts have acquired immunity via multiple matches will tend to go extinct earlier than those with fewer matches (Fig. 3b, Fig. S6). Somewhat paradoxically, this orderly extinction will facilitate in turn a new viral escape and the stochastic initiation of another virus expansion cycle, as it reduces nestedness by preferentially removing viruses with a high number of matches. Hence, resistance to viruses declines at the population level, rather than at the individual host level. Specifically, the extinction process increasingly focuses the competition for hosts between viruses of two kinds, those that have either no match (0-match) and those with a single match (1-match) (Fig. 4b). Their relative abundances increase in the population, with an increase in the proportion of 1-matches raising the effective mutation rate of the overall remaining viral population (Supplementary Methods). Viral escapes are indeed associated with a particular tripartite structure in which hosts are immune to viruses via a single match (Fig. S7).

**Fig. 4.**
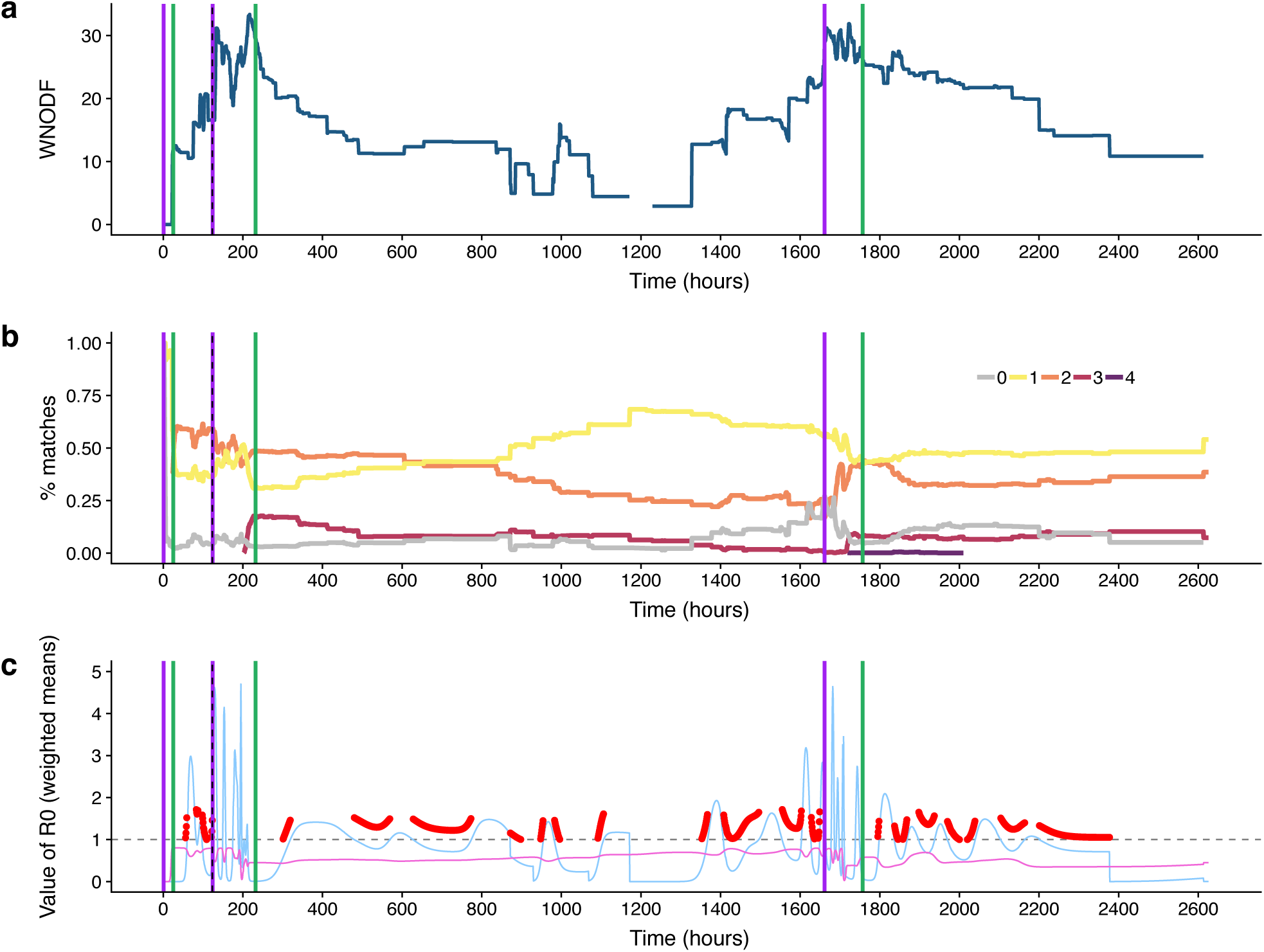
Processes leading to regime shifts. **a**, Weighted nestedness increases during VDRs and decreases during HCRs. **b**, The disassembly of the network during HCRs increases the proportion of 0- and 1-matches, particularly towards the end of these periods. **c**, During HCRs the average reproductive number 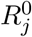 (light blue line) is approximately 1 (horizontal dashed line). Although the mean 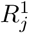 is always < 1 (pink), *R*_*pot*_ can be > 1. During these times, depicted with red points, a mutation would allow the viral population to grow. Because mutations are more likely to occur in highly abundant virus strains, it becomes likely that a mutation hits a virus with a high potential for virus growth, and therefore, escape.

### Epidemiologically, virus escape depends on 0 and 1 CRISPR matches

Epidemiologically, these two processes, viral growth rate, and potential escape, can be captured via two modified measures of the basic reproductive number^27^ (Supplementary Methods). The first, 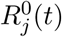, is the number of offspring produced at time *t* by the virus strain *j* from infecting all hosts with no protection to it (0-matches). The second, 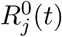, is the number of offspring that a virus strain *j* would produce by escaping protection from hosts via a single mutation (1-match) at time *t*. These two components can be added to obtain the ‘potential reproductive number’ of a virus strain *j*, 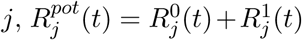, which quantifies the contribution of a viral strain and its potential progeny to the population growth, conditional on escape. During HCRs, the contribution of the second component 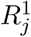 can raise 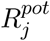 above the critical value of 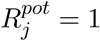 (Fig. 4c), indicating a viable escape. Towards the end of an HCR, small viral outbreaks (Fig. 1b) locally generate some virus diversification which counterbalances extinctions. Extinctions and births of virus strains change both the size of the immunity network and the distribution of protection weights among the links. These two effects are reflected in the nestedness index^26,28,29^.

### Results hold for multiple simulations

To ensure the generality of our results given the stochastic nature of the dynamics, we repeated our analysis for 100 simulations. Across these simulations, we find the same characteristic differences between the regimes in dynamics and network structure as in our main example. In particular, VDRs are shorter than HCRs, have higher values of weighted nestedness, have more modules in the infection networks, and have higher values of 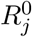 (Fig. S8). Together, these results indicate a general pattern of viral diversification via the creation of host niches, the corresponding buildup of immunity in VDRs, and the breakdown of weighted nestedness that eventually leads to escape or extinction during HCRs.

### Empirical immunity networks are weighted-nested

Finally, we analyzed the structure of immunity in three empirical systems because the weighted nestedness of the protection network is a key structural feature in the shifts between dynamical regimes. Ideally, one would consider temporal data on host-virus CRISPR matches. Such data are currently unavailable as obtaining it requires hundreds of genotypes of virus and CRISPR alleles to be resolved. Metagenomic datasets^21^ can obtain such diversity but do not link spacers to individual host strains. Hence, currently available temporal data are either resolved at the strain level but for very low numbers of strains^4,22^ or contain many spacers and protospacers but without host-virus CRISPR matches between multiple strains^21^. In fact, even for non-temporal data, there are very few examples of virus and host populations sufficiently resolved at an individual strain level. We considered here three such static data sets, and asked whether they exhibit evidence for the weighted-nested immunity structure predicted by the theory.

In our empirical data sets both CRISPR alleles and virus strains have been carefully and manually assembled, and we had previously established their profiles of diversity^24,30,31^ but not their structure. The data represent lytic, chronic, and temperate virus lifestyles and two different microorganisms from the two domains of life where CRISPR occurs (Archaea and Bacteria). The first two data sets compare genomes resolved to individual strains of the thermoacidophilic crenarchaeon *Sulfolobus islandicus* sampled at a single time point from two different locations. The third data set is from the Gammaproteobacteria *Pseudomonas aeruginosa*^31^, isolated from the sputum of CF patients in a high resolution longitudinal dataset collected by Marvig et al.^32^, with virus genomes obtained from the temperate mu-like viruses from the genomic database because these are the most highly viral genomes in *P. aeruginosa*.

Because we did not have a time series of matches, our analysis cannot directly address transitions between HCR and VDR regimes. The data does allow us to interrogate the structure of immunity about the existence of weighted nestedness, a key feature indicative of such dynamics. From each data set, the empirical immunity matrices were constructed by comparing CRISPR arrays from each individual host strain to virus genomes. We assessed the statistical significance of nestedness for each empirical network by comparing the observed value of its nestedness index (for two different indices, see Methods) to a distribution obtained from 10,000 shuffled networks. We find that the probability that a shuffled network will be more nested than the observed one is effectively nil (p-value< 0.0001; Methods) in all three empirical networks (Fig. 5).

**Fig. 5.**
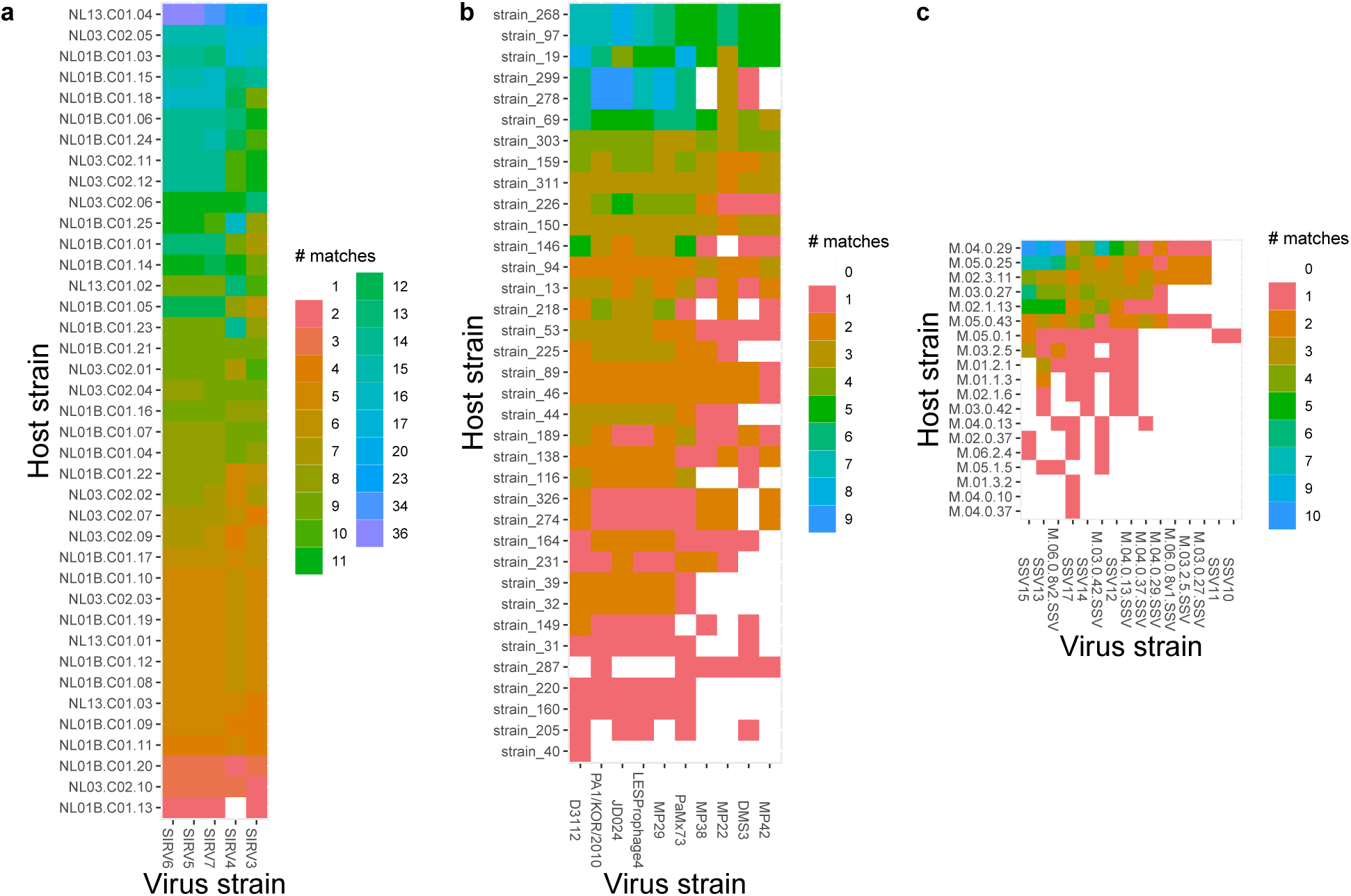
Weighted nestedness of empirical immunity networks. The matrices depict the number of shared spacers and protospacers between hosts (rows) and viruses (columns). Each network is ordered by column and row sums and is nested in the quantity of matrix cells. The networks come from 3 different systems: **a**, *Sulfolobus islandicus* hosts from a single location in Yellowstone National Park compared to contemporary lytic SIRV viruses isolated from Yellowstone National Park. **b**, *S. islandicus* hosts compared to contemporary chronic SSV viruses from the Mutnovsky Volcano in Russia, 2010. (**c**, *Pseudomonas aeruginosa* hosts from Copenhagen compared to temperate mu-like viruses. To test the hypothesis that for a given distribution of matches in the population of host strains, the observed network is organized in a non-random, weighted-nested pattern, we shuffled networks by randomly distributing the interactions.

In light of our theoretical results, these findings suggest that in these three systems, hosts may control populations of viruses via distributed immunity that is redundant in the number of matches. Moreover, we find that, as in our theoretical results, the empirical host-spacer networks in the three VDRs are also significantly modular (Fig. S9), with only one out of 5 modules showing a phylogenetic signal (Table S3). This suggests that the mechanism by which immunity is obtained is similar to the one we have described: modules delineate groups of host strains that share immunity via the same spacers, and are therefore susceptible to similar viruses.

## Discussion

The role of resistance and immunity in driving the diversity, evolution, and coevolution of microbes and viruses has been extensively studied^2,4,21,33–37^. The role it plays in structuring microbial populations remains however poorly investigated, despite possible consequences for host-virus coevolution and population stability. Our theoretical results show that the coevolutionary dynamics of a host-pathogen system with CRISPR-induced resistance can influence, and in turn be influenced by, the network structure of strain diversity. In particular, modularity of the infection network and weighted nestedness of the immunity network interact to affect the transient nature of alternating dynamical regimes. By promoting pathogen diversification, the former builds the latter. In turn, the weighted nestedness puts an end temporarily to pathogen diversification as it implies redundant resistance that virus mutation can no longer overcome. It also contains the seeds of its own unraveling by implying a certain order in viral extinction, and in so doing, creating the conditions for a potential new viral escape. Thus, the weighted nestedness of protection networks is at the center of the coevolutionary dynamics and the transient dynamical regimes we have described here.

Modularity has been described for host-parasite infection networks and attributed to a variety of mechanisms such as phylogeny, specificity and coevolution^9,14,38^. It was also detected in a host-virus system, where it was suggested to arise via negative frequency-dependent selection^16^. Its emergence from immune selection was demonstrated here. It is consistent with existing strain theory developed mostly for human infections and pathogens with multilocus encoding of antigens, when evolution is considered explicitly^17,39–42^. Strain theory posits that immune selection acts as a form of negative frequency-dependent selection, creating groups of pathogens with limiting overlap in antigenic space. Pathogens with antigens that are novel to the host immune system are at a competitive advantage, while those with common ones are at a disadvantage. In the absence of data on immunity of hosts, this pattern has been described as modules, or clusters, in networks that depict genetic similarity between the pathogens themselves^17,18^. In contrast to strain theory, we are able to consider here the immune genotype of pathogens and hosts simultaneously, which allows us to move beyond pathogen genetic similarity to consider actual infection patterns. Here, niche structure arises in part from the addition of new spacers along with competition among viruses for hosts.

One major difference with previous strain theory is that modularity and associated coexistence in our model are only transient and not accompanied by stable population dynamics. Future work should examine whether this difference and the associated shifts in dynamical regimes arise from the heritable nature of CRISPR-based immunity, which by definition cannot produce fully susceptible offspring, needed to sustain transmission. Host-virus coexistence can also result from loss of immunity^36^, a feature absent from our model and that could prolong VDRs by supplying a constant inflow of susceptible hosts.

We show that, when multiple viruses are involved, weighted nestedness with multiple spacers derived from repeated infections provides a population-level mechanism by which hosts can control viral populations. It remains an open question whether it can arise under non-CRISPR immunity or non-heritable immunity. For CRISPR-induced immunity, nestedness reflects the coevolutionary diversification of hosts in response to that of viruses, and more specifically, the resulting redundancy of immune memory, which allows hosts to keep viruses in check despite their occasional escape. Thus, observation of weighted nestedness in nature would be an indication of such control. Variation of this network property over time contains valuable information on the relative pace of viral diversification and host acquisition of immunity. Observing the temporal course of nestedness in both dedicated experiments and in natural environments should shed light on its role for controlling virus populations in relation to host and virus diversification. Quantitative extensions should also investigate appropriate ways to normalize weighted nestedness (sensu Song et al.^29^), so that effects of its change can be isolated for network size and distributed redundancy of protection.

The transience of alternating dynamical regimes in the CRISPR system is not an impediment to the formation of rich structures by coevolution; it is here its natural consequence. It should be recognized, however, that the alternation of dynamical regimes occurs in the model in a defined region of parameter space that allows for high diversification, and is absent in some earlier simulations (e.g. Childs et al.^25^). Because host-control periods lead to stochastic viral extinction, identifying the conditions in terms of critical parameter combinations that give rise to these dynamics is of practical importance.

Future work in this area should consider the role of demographic stochasticity arising from small population numbers and discrete individuals. Our numerical implementation of the system treated population dynamics as deterministic, and therefore relied on a given abundance threshold for extinction. It also initialized the size of a new strain at a given abundance slightly above threshold. We expect our results to hold qualitatively even though there should be some quantitative differences in the exact parameter values for which the two implementations are comparable. This should be the case for the mutation rate as the emergence probability of new strains should decrease due to demographic stochasticity. For a fully stochastic implementation, the analytical expressions of Chabas et al.^27^ could be extended to compute and understand emergence probability under varying frequencies of resistance alleles. A future theory should also more broadly investigate the dynamics-structure nexus over parameter space, in particular for protospacer mutation rates and spacer acquisition probabilities, which set the pace of evolution and have already been recognized as key to the outcome of viral extinction^22,25,43^. Finally, anti-CRISPR mechanisms maintained by viruses may also affect network structure.

The demonstration of significantly weighted nestedness in the empirical networks suggests that coevolution involving CRISPR-induced immunity is at play in natural microbial populations. It further suggests the applicability of our modeling results to predict these kinds of transitions. Obtaining time series data on CRISPR-mediated interactions between multiple host and virus strains should be a priority to further test our theoretical results.

Ultimately, knowledge on the stability of diverse host and viral populations can be applied to control of microbes in food and industrial sciences, infectious disease emergence and treatment with phage therapy, microbiome dynamics, agriculture, and environmental engineering.

## Data and code availability

Simulated and empirical network data, as well as code for simulations and their analysis are available in the dedicated GitHub repository associated with this paper at: https://github.com/EcologicalComplexity-Lab/CRISPR_networks.

## Methods

### The model

The model implements the formulation by Childs et al.^20^, which combines three main components: ecological population dynamics, stochastic coevolution generating diversity, and molecular identity of hosts and viruses defining CRISPR immunity. Diversification events (spacer acquisition by the host and protospacer mutation of viruses) are modeled stochastically, whereas population dynamics of different sub-populations (i.e., strains) of hosts and viruses are represented deterministically with Lotka-Volterra differential equations (eqns. 1,2).

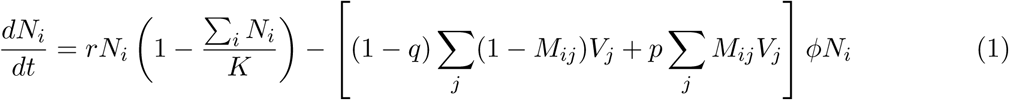

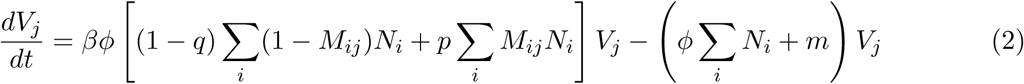

Each strain of host *i* and virus *j* is defined by a *unique* genomic state of their spacer and protospacer sets *S*_*i*_ and *G*_*j*_, respectively. In the ecological population dynamics, each host strain *i* has abundance *N*_*i*_ (the carrying capacity of all strains is *K*) and reproduces at a per-capita rate of *r*. Each viral strain *j* has abundance *V*_*j*_, which increases due to infection and lysis of hosts and decays at a density-independent rate *m*. Extinction of any host or viral strain occurs when these fall below a critical threshold *ρ*_*c*_. Viruses infect at a constant ‘adsorption rate’ *ϕ* either hosts that do not have protection or those whose protection fails (see below).

Host immunity to a virus is defined in the molecular component of the model and is based on genomic sequence matches between the spacer and protospacer sets. Specifically, CRISPR immunity is defined using the function *M*_*ij*_ = *M* (*S*_*i*_, *G*_*j*_), which equals 1 if there is at least 1 match between the sets: |*S*_*i*_ ∩ *G*_*j*_| ≥ 1, or 0 otherwise. The CRISPR immune mechanism is not perfect and can fail. When *M*_*ij*_ = 1, there is a probability *p* that the host strain is lysed and correspondingly, 1 − *p* that it survives and the virus is eliminated. On the other hand, when *M*_*ij*_ = 0, there is a probability *q* that the virus strain is eliminated, resulting in the acquisition of a protospacer by the host, and 1 − *q* that it is lysed by the virus. Both *p* and *q* are small (*p, q* ≪ 1).

The above Lotka-Volterra dynamics are implemented in between any two stochastic events concerning evolution. Specifically, errors in viral replication can result after successful infection of a host, leading to the replacement of a random protospacer with mutation rate *µ* (per protospacer per viral replication). This incorporates a new viral strain into the system with an initial low abundance of 1.1 times the extinction threshold *ρ*_*c*_. Viruses cannot mutate multiple protospacers at the same time. During an unsuccessful infection attempt by a virus (regardless of host immunity), there is also a probability *q* of acquiring a new spacer by incorporating a protospacer and integrating it into the host’s CRISPR system at its leading end. Hosts cannot acquire multiple spacers at the same time. The maximum number of spacers per hosts strain is constant. If the maximum number of acquired spacers is reached, the addition of a spacer to the leading end is accompanied by the deletion of a spacer at the trailing end. In our simulations, the length of the spacer cassette is set to a sufficiently large value to avoid loss of acquired immune memory.

We numerically implemented the model in *C++* to increase computational efficiency. This enables consideration of a larger number of spacers/protospacers, and hence host and virus richness, for longer simulation times, than in the original MATLAB code^20^). Our implementation combines: (1) a Gillespie algorithm to determine the time between two stochastic events and to randomly select a virus/host strain to mutate a protospacer/acquire a new spacer, and (2) a numerical ODE solver using Euler’s method. The code includes the following features to facilitate subsequent network (and other) analyses: (1) all data related to virus or host strains (e.g., identity of protospacers/spacers, abundance values) at each time are written to files during simulation; (2) the parent ID of each newly generated virus or host strain is tracked to generate phylogenetic trees; (3) a checkpoint implementation for running longer simulations. Details on the implementation are in Supplementary Information.

We used the parameters summarized in Table S4 based on^20,25^. Host growth rate corresponds to a doubling time of an hour within the range of observed values for *P. aeruginosa* (20 mins) and *Sulfolobus* (8 hrs). The adsorption rate also falls within typical range values of 10^−8^ to 10^−9^ ml min^−1^. The order of magnitude of our mutation rate is consistent with standard mutation rates for DNA based organisms, if we assume that mutation at a single site enables protospacer escape from a spacer match. The burst size of 50 is on the low side with values of 200 observed in *P. aeruginosa*. This is motivated by a lower protospacer number than currently documented in nature. Specifically, the speed of virus escape ultimately depends on the effective rate of mutation per protospacer, which increases linearly with the product of burst size and the mutation rate parameter *µ* and decreases inversely with the number of protospacers. Therefore, our parameter set should generate similar speeds of protospacer evolution than for higher burst sizes and also higher numbers of protospacers (e.g. *β* = 200 and *g*_*p*_ *×* 4). Our numerical implementation allows consideration of a higher number of protospacers than before; for simplicity, in our study we used a number of protospacers that is still below those typically observed but have compensated with a lower value of offspring produced (burst size; see SI.)

### Definition of regimes

Our general approach to define regimes was to classify each point in the virus abundance time series to either a HCR or a VDR. We did so by detecting changes in relative virus abundance, defined as the total virus abundance at any given time *t, V*_*T*_ (*t*), divided by the maximum abundance in whole time series, *V*_*T*_ : *A*(*t*) = *V*_*T*_ (*t*)*/max*(*V*_*T*_). Using relative abundance allows for comparisons and analysis across multiple simulations; considering absolute abundance does not change the regime definition (Fig. S17). A HCR is a sequence of points in which *V*_*T*_ (*t*) changes very little, which can be captured by calculating the changes between consecutive time points. We consider the first difference *A*′ = *A*(*t*_*x*_) − *A*(*t*_*x*−1_), and the second difference *A*′ = *A*′(*t*_*x*_) − *A*′(*t*_*x*−1_). Values close to 0 in *A*′ would occur when (1) there is no, or low, change in A between two consecutive time intervals so that A’ itself is small in each of these periods; or (2) *A*′ varies and changes sign in these consecutive intervals but the size of these changes ‘up’ and ‘down’ are small, as well as the peak in *A* this reflects. (We ignore here the case of an almost linear increase, which does not apply). We classified each point as belonging to the HCR if the absolute value of *A*′ is smaller than a given small threshold, so that 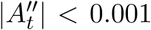, and to a VDR otherwise. This procedure creates a sequence of points *C*(*t*) with each point classified into HCR or VDR (Fig. S10). We added two further conditions. The first one includes time points in the HCR when isolated increases in virus abundance are not recognized as sufficiently small peaks by our threshold, but these increases are localized outbreaks in the sense of not esulting in a complete escape and a regime switch. To include such events within the HCR we calculated a threshold *f* defined as the 75% quantile value of the distribution of VDR lengths in *C*. Any sequence of VDR points shorter than *f* was converted to the HCR. The last condition for classification to a HCR was that a sequence of points had to be longer than the longest-lasting virus outbreak.

While the values for the thresholds we have used may seem, to some degree, arbitrary, this would be the case for any other algorithm for classification because a ‘regime’ is not naturally defined by the dynamical system, but rather detected in the emerging time series. As an independent corroboration for our method, we imposed the regime boundaries calculated using virus abundance time series to time series of host abundance and virus diversification. The regimes are evident in those time series as well (e.g., Fig. S1). For example, in the virus diversification time series, the rate of diversification is much steeper in the VDR than in the HCR (Fig. S3). This additional verification increases the confidence we have in our ability to classify these regimes.

### Network construction

The networks we use are bipartite networks, which contain two sets of nodes (e.g., hosts and spacers or hosts and viruses). Bipartite networks are mathematically represented using incidence matrices. A graphical overview of the networks (and associated matrices) and how we constructed them can be found in Fig. 2. Here, we provide details on network construction.

### Networks of genetic composition

At each time *t*, we defined a *host-spacer network*, 𝒮(*t*) (and associated matrix **S**_*xi*_(*t*)) as a bipartite network in which each edge is drawn between a host strain *i* and a spacer *x* (Fig. 2a,e). We analogously defined a *virus-protospacer network* 𝒫(*t*) (and associated matrix **P**_*yj*_(*t*)) as a bipartite network in which each edge is drawn between a virus strain *j* and a protospacer *y* (Fig. 2b,f). Hence, in each network a host or virus strain’s genome is the set of its neighboring spacer or protospacer nodes, respectively.

### Immunity network

We used 𝒮(*t*) and 𝒫(*t*) to define an *immunity network* at a time *t*, ℐ(*t*) (and associated matrix **I**_*ij*_(*t*)) in which edges are drawn between a virus strain *j* and a host strain *i*. Edge weights were defined as the number of matching spacers and protospacers between 𝒮(*t*) and 𝒫(*t*) for any given host-virus pair (Fig. 2c,g). That is: 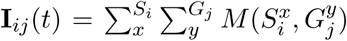, where *M* has a value of 1 if spacer *x* in the spacer set of host *i, S*_*i*_, is the same as protospacer *y* in the protospacer set of virus *j, G*_*j*_ and 0 otherwise.

### Infection network

We defined an *infection network* at a time *t*, 𝒢(*t*) (and associated matrix **G**_*ij*_ (*t*)) as a subset of the immunity network in which **I**_*ij*_ (*t*) = 0 (Fig. 2d,h). That is, edges in **G**_*ij*_ (*t*) were drawn between hosts and viruses that did *not* have an edge in ℐ. We weighted the edges of **G**_*ij*_ (*t*) by a normalized measure of encounter between virus strain *j* will and a host *i*, based on their abundance values: 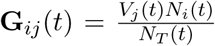, where *V*_*j*_ (*t*) and *N*_*i*_(*t*) are the abundances of a virus strain *j* and a host strain *i* at time *t* and *N*_*T*_ is the total abundance of hosts.

### Network analysis

#### Weighted nestedness

We evaluated the nestedness of **I**_*ij*_ (*t*) using WNODF^26^. Briefly, the index ranges from 0 to 100, with the maximum 100 representing perfect nestedness. Perfect nestedness occurs when all 2×2 sub-matrices of the form

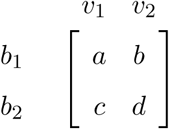

satisfy the conditions for host *b*_1_ to be immune to the two viruses via more matches than *b*_2_ (*a* > *c, b* > *d, a* > *d*) and for *v*_2_ to have less matches to the two hosts than *v*_1_ (*a* > *b, c* > *d*). We calculated WNODF with the `networklevel` function in the bipartite package (version 2.11) in R.

We calculated WNODF when at least 2 strains of both hosts and viruses were present.

In empirical data we evaluated nestedness in two ways. First, we used WNODF. Second, we calculated the largest eigenvalue of 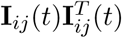, *ρ*, as suggested by^44^. We did not use *ρ* in the simulations because preliminary analyses indicated that this measure is highly dependent on network size and therefore cannot be used to compare between networks. However, it is suitable for comparing an observed network to its shuffled counterparts because for a given distribution of weights an network size high values of *ρ* indicate a more quantitatively nested structure^44^. WNODF and *ρ* are the only two measures for quantitative nestedness we are aware of. WNODF cannot handle networks which are fully connected (i.e., density of 1) and we therefore did not use it for our Yellowstone data set.

#### Community detection

To find ‘communities’, or as commonly referred to, ‘modules’, we used the map equation objective function to calculate the optimal partition of the network^45,46^ with the R package infomapecology (version 0.1.2)^47^. Briefly, the map equation is a flow-based and information-theoretic method (implemented with Infomap), which calculates network partitioning based on the movement of a random walker on the network (see for details). In any given partition of the network, the random walker moves across nodes in proportion to the direction and weight of the edges. Hence, it will tend to stay longer in dense areas representing groups of, for example, viruses and hosts with high interaction density. These areas can be defined as ‘modules’. The time spent in each module can be converted to an information-theoretic currency using an objective function called the map equation and the ‘best’ network partition corresponds is the one that minimizes the Map Equation^45,46^. For convenience we use the term ‘modules’, as it is commonly used to refer partitions of networks across different disciplines, but Infomap does not calculate a modularity function (sensu^48^). We chose the map equation over the more commonly used modularity objective function because of the computational efficiency of its implementation (Infomap)^49^, given that we needed to analyze hundreds of thousands of networks.

#### Significance of modularity and weighted nestedness

To test if a given network structure is non-random, we evaluated the statistical significance of the structural index (map equation value for modularity and WNODF or *ρ* for weighted nestedness) by comparing it to a distribution of the index for shuffled versions of the matrix. We shuffled binary matrices (host-spacer and infection networks) with function `r00` in package `vegan` (version 2.5-4) in R), an algorithm that maintains the density of the network. This was done internally using infomapecology (version 0.1.2)^47^. We shuffled weighted matrices (immunity networks in simulated and empirical data) by randomly distributing the interactions (function `r00_samp` in package `vegan` (version 2.5-4) in R). This algorithm maintains the density of the network and the distribution of weights while shuffling the structure. Therefore, it tests the hypothesis that the for a given distribution of matches the observed network is non-randomly structured in a quantitatively nested way.

### Phylogenetic signal in modules

We first computed the pairwise distances between pairs of strains within each module from branch lengths, with function `cophenetic.phylo` in package ape in R. Then, we permuted the identity of strains in modules, maintaining module size, and recalculated the mean pairwise phylogenetic distance. The null hypothesis is that the permuted distance is smaller than the observed one (per module), and therefore there is no phylogenetic signal. Rejecting this hypothesis indicates a phylogenetic signal because the observed phylogenetic distance between hosts within each module would be smaller than expected by chance (closely related hosts share a module). Because each network has several modules, the threshold for significance was adjusted by dividing 0.05 by the number of modules (Bonferroni correction).

### Epidemiological measures

We derived two modified measures of the basic reproductive number (see derivation in Supplementary Methods). The first, 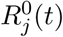, is the number of offspring produced at time *t* by virus strain *j* from infecting all hosts with no protection to it (0-matches) and is defined as:

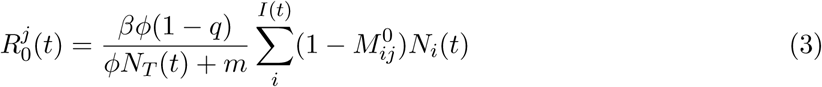

The delta-function 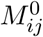 equals 1 (and 0 otherwise) when there are no matches between the set of spacers of host *i, S*_*i*_ and the set of protospacers of virus *j, G*_*j*_. *I*(*t*) is the set of hosts in time The second, 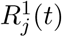, is the number of offspring that a virus strain *j* would produce by escaping protection from hosts via a single mutation (1-match) at time *t* and is defined as:

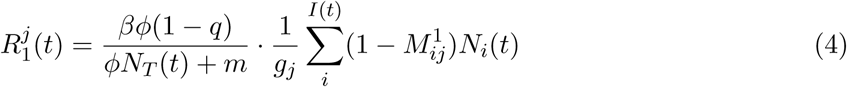

Here, 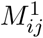 equals 1 (and 0 otherwise) when there is exactly 1 matching spacer-protospacer pair in the set of spacers of host *i, S*_*i*_ and the set of protospacers of virus *j, G*_*j*_. *g*_*j*_ is the length of the protospacer cassette of virus *j* and quantifies the probability that a mutation will hit a particular protospacer. Adding these two quantitites together we obtain the ‘potential reproductive number’ of a virus strain *j*, which quantifies the contribution of a viral strain and its potential progeny to population growth, conditional on escape:

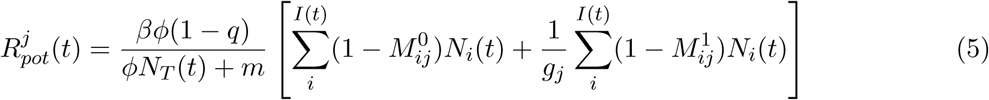

### Empirical data

The first data set represents a single time point from three adjacent hot springs (NL01, NL10 and NL13) in the Nymph Lake region of Yellowstone National Park. These three springs have shown to share a single well-mixed population of *S. islandicus hosts*^50^. Viruses were collected from contemporary populations and CRISPR spacers and virus isolates suggest that these are dominated by the lytic virus SIRV^51^. The second data set is from *S. islandicus* strains isolated from several springs in the Mutnovsky Volcano in Kamchatka Russia that also were shown to share populations of hosts. The predominant viral population in these hosts is the chronic Sulfolobus Spindle Shaped Virus (SSV)^24^. The third data set consists of longitudinal sampling of human-adapted *P. aeruginosa* isolates from sputum samples of Cystic Fibrosis patients collected at a hospital in Copenhagen, Denmark and a global set of temperate mu-like viruses from *P. aeruginosa* viruses extracted from NCBI to substitute for a lack of sequenced contemporary viruses^31,32^. See detailed methods in Supplementary Methods.

## Acknowledgments

We thank Whitney England and Matthew Pauly for data processing and collection, and Qixin He for guidance on the construction of the phylogenetic trees from model outputs. RW acknowledges the support of the Cystic Fibrosis Foundation (CFF C2480) and an Allen Distinguished Investigator Award from the Allen Frontiers Institute; MP, that of the University of Chicago. We are also grateful for the access to the computer cluster of the Research Computing Center (RCC) of the University of Chicago.

---

**Table S1.** Lack of phylogenetic signal in modules in a simulated infection network. The first VDR contained too little hosts and viruses to analyze.

| VDR ID | Module | Size | Pvalue | Signal |
| --- | --- | --- | --- | --- |
| 2 | 1 | 2 | 0.61 | No signal |
| 2 | 2 | 2 | 0.93 | No signal |
| 2 | 3 | 30 | 0.10 | No signal |
| 2 | 4 | 12 | 0.01 | Signal |
| 2 | 5 | 6 | 0.46 | No signal |
| 2 | 6 | 6 | 0.67 | No signal |
| 2 | 7 | 12 | 0.23 | No signal |
| 2 | 8 | 12 | 0.99 | No signal |
| 2 | 9 | 20 | 0.27 | No signal |
| 2 | 11 | 12 | 0.80 | No signal |
| 2 | 12 | 6 | 0.50 | No signal |
| 2 | 13 | 2 | 0.24 | No signal |
| 2 | 14 | 6 | 0.15 | No signal |
| 3 | 1 | 90 | 0.40 | No signal |
| 3 | 2 | 56 | 0.65 | No signal |
| 3 | 3 | 72 | 0.63 | No signal |
| 3 | 4 | 12 | 0.11 | No signal |
| 3 | 5 | 72 | 0.00 | Signal |
| 3 | 6 | 30 | 0.32 | No signal |
| 3 | 7 | 6 | 0.54 | No signal |
| 3 | 8 | 6 | 0.05 | No signal |
| 3 | 9 | 6 | 0.71 | No signal |
| 3 | 10 | 2 | 0.79 | No signal |

**Table S2.** Lack of phylogenetic signal in modules in a simulated host-spacer network.

| VDR ID | Module | Size | Pvalue | Signal |
| --- | --- | --- | --- | --- |
| 1 | 1 | 6 | 0.38 | No signal |
| 1 | 2 | 12 | 0.41 | No signal |
| 1 | 3 | 6 | 0.10 | No signal |
| 1 | 4 | 2 | 0.91 | No signal |
| 2 | 1 | 462 | 0.58 | No signal |
| 2 | 2 | 156 | 0.08 | No signal |
| 2 | 3 | 182 | 0.47 | No signal |
| 2 | 4 | 156 | 0.06 | No signal |
| 2 | 5 | 132 | 0.64 | No signal |
| 2 | 6 | 110 | 0.47 | No signal |
| 2 | 7 | 90 | 0.86 | No signal |
| 2 | 8 | 72 | 0.37 | No signal |
| 2 | 9 | 72 | 0.15 | No signal |
| 2 | 10 | 72 | 0.37 | No signal |
| 2 | 11 | 42 | 0.95 | No signal |
| 2 | 12 | 42 | 1.00 | No signal |
| 2 | 13 | 20 | 0.29 | No signal |
| 2 | 14 | 6 | 0.08 | No signal |
| 2 | 15 | 2 | 0.38 | No signal |
| 2 | 16 | 2 | 0.16 | No signal |
| 3 | 1 | 1056 | 0.02 | Signal |
| 3 | 2 | 992 | 0.04 | Signal |
| 3 | 3 | 1560 | 0.00 | Signal |
| 3 | 4 | 992 | 0.00 | Signal |
| 3 | 5 | 462 | 0.00 | Signal |
| 3 | 6 | 110 | 0.00 | Signal |
| 3 | 7 | 182 | 0.03 | Signal |
| 3 | 8 | 72 | 0.13 | No signal |
| 3 | 9 | 42 | 0.92 | No signal |
| 3 | 10 | 20 | 0.45 | No signal |
| 3 | 11 | 30 | 0.01 | Signal |
| 3 | 12 | 12 | 0.67 | No signal |
| 3 | 13 | 2 | 0.94 | No signal |
| 3 | 14 | 2 | 0.07 | No signal |
| 3 | 15 | 2 | 0.59 | No signal |
| 3 | 16 | 2 | 0.07 | No signal |

**Table S3.** Lack of phylogenetic signal in modules in an empirical host-spacer network for data set Russia 2010.

| Module | Module size | Signal |
| --- | --- | --- |
| 1 | 2 | No signal |
| 4 | 2 | No signal |
| 5 | 12 | No signal |
| 6 | 2 | Signal |
| 11 | 2 | No signal |

**Table S4.** Description and values of model parameters used in simulations, following Childs, L. M., Held, N. L., Young, M. J., Whitaker, R. J. & Weitz, J. S. Multiscale model of CRISPR-induced coevolutionary dynamics: diversification at the interface of Lamarck and Darwin. Evolution 66, 2015–2029 (2012).

| Parameters | Description | Value |
| --- | --- | --- |
| $r$ | Growth rate | $1h^{-1}$ |
| $K$ | Carrying Capacity | $10^{5.5} \text{ mL}^{-1}$ |
| $\beta$ | Burst size | 50 |
| $\phi$ | Adsorption rate | $10^{-7} \text{ mL/h}$ |
| $m$ | Viral decay rate | $0.1 \text{ h}^{-1}$ |
| $\mu$ | Viral mutation rate per protospacer and replication | $1 - 5 \times 10^{-7}$ |
| $\rho_c$ | Density cutoff | $0.1 \text{ mL}^{-1}$ |
| $p$ | CRISPR failure probability per infection | $10^{-5}$ |
| $q$ | Spacer acquisition probability per infection | $10^{-5}$ |
| $g_p$ | Number of protospacers in a virus strain | 15 |
| $g_s$ | Number of spacers in a host strain | 10 |

**Fig. S1.**
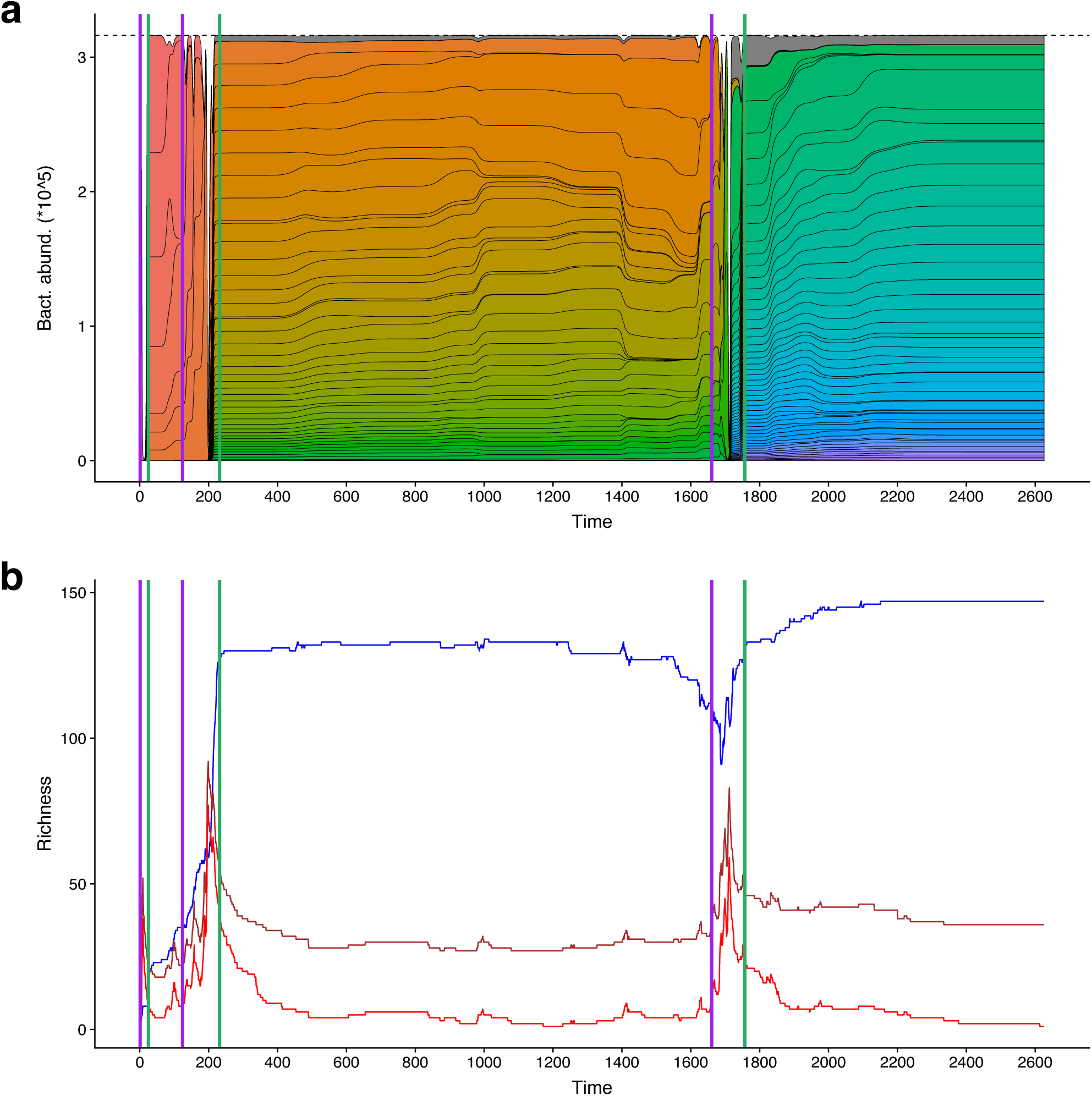
Viral and host abundance and richness. **a** Host abundance. The 100 most abundant strains are colored, the rest are aggregated and shown in gray. **b** Richness (i.e., number of unique strains) of hosts (blue) and viruses (red). The number of unique spacers (spacer richness) is depicted in brown. During VDRs the abundance of both hosts and viruses fluctuates. As a response to virus diversification, host richness eventually increases despite possible declines at the beginning of the VDR resulting from the initial viral attack.

**Fig. S2.**
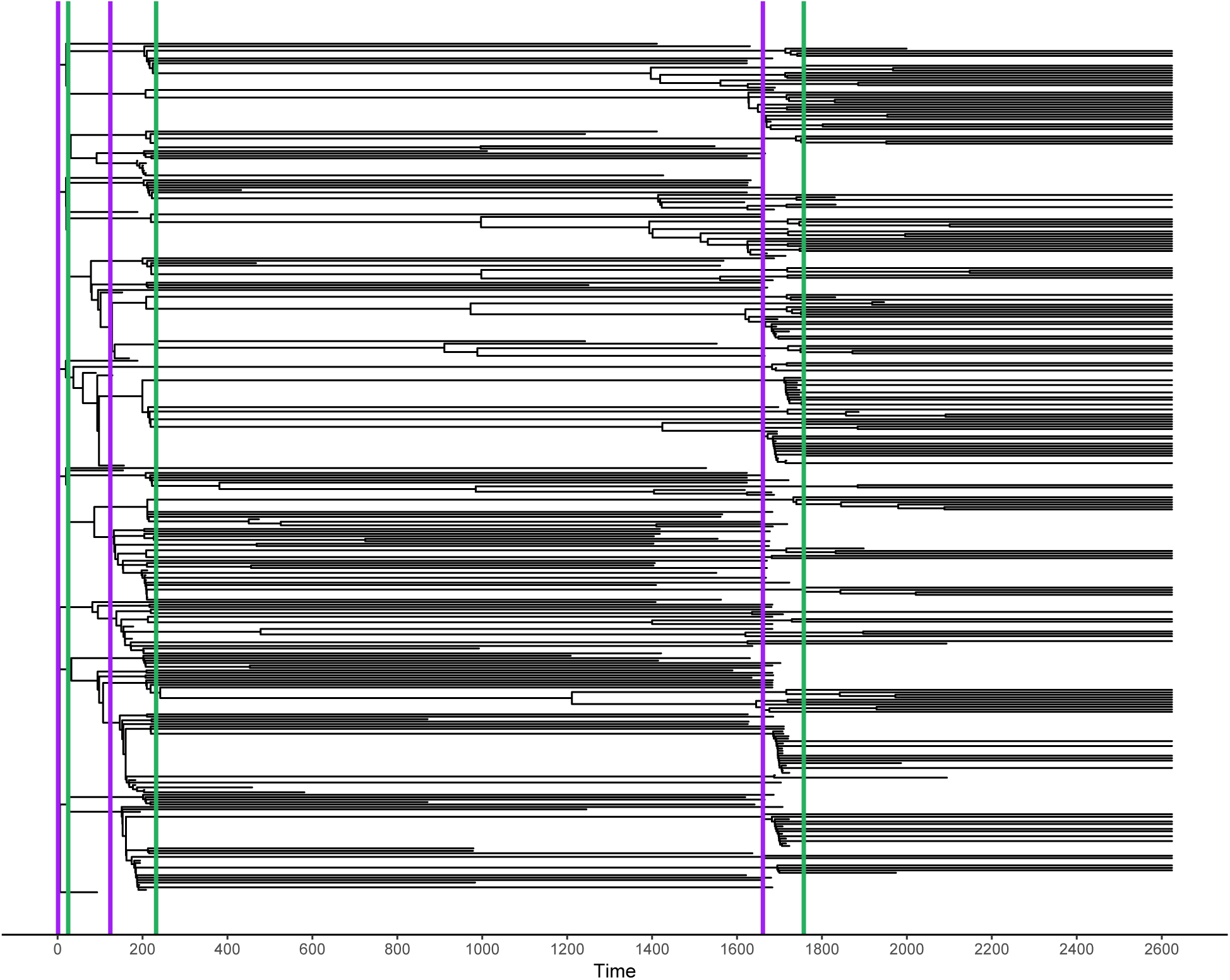
Host phylogenetic tree. The tree is not inferred, but rather drawn based on exact genealogical data (which strains descends from which) collected during the simulation. Branch length indicates the lifetime of any given host strain. Hosts diversify and go extinct primarily during VDRs, resulting in strain replacement but strains can also persist from one VDR to the next.

**Fig. S3.**
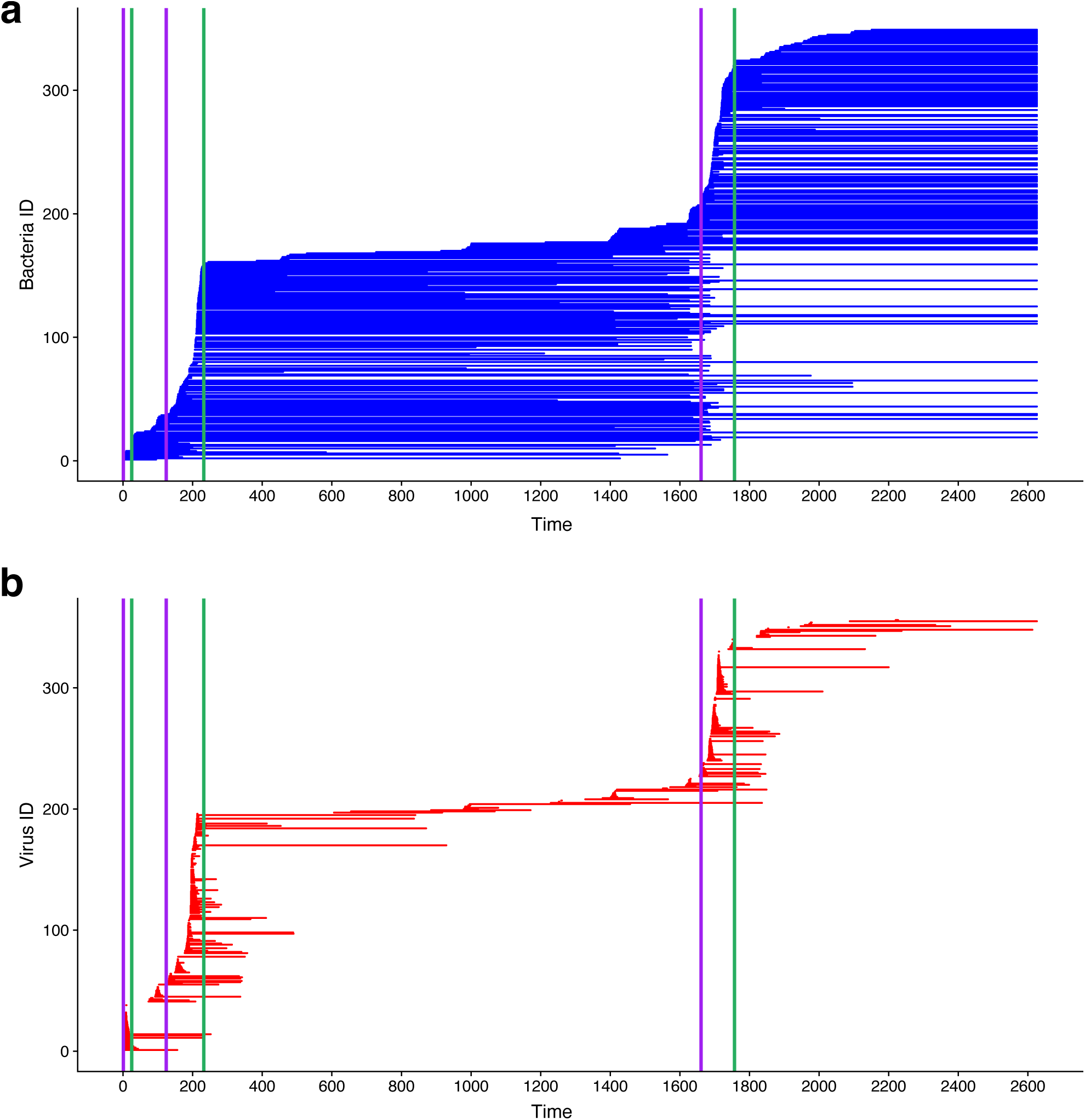
Host and virus diversification and extinction. Each host **a** or viral **b** strain is plotted with a line, starting at the time when the strain was generated and ending when the strain went extinct. During VDRs the rate of diversification of both viruses and hosts is higher than during HCRs. While viruses have relatively short persistence (shorter line lengths), hosts have long persistence and can persist during an entire VDR.

**Fig. S4.**
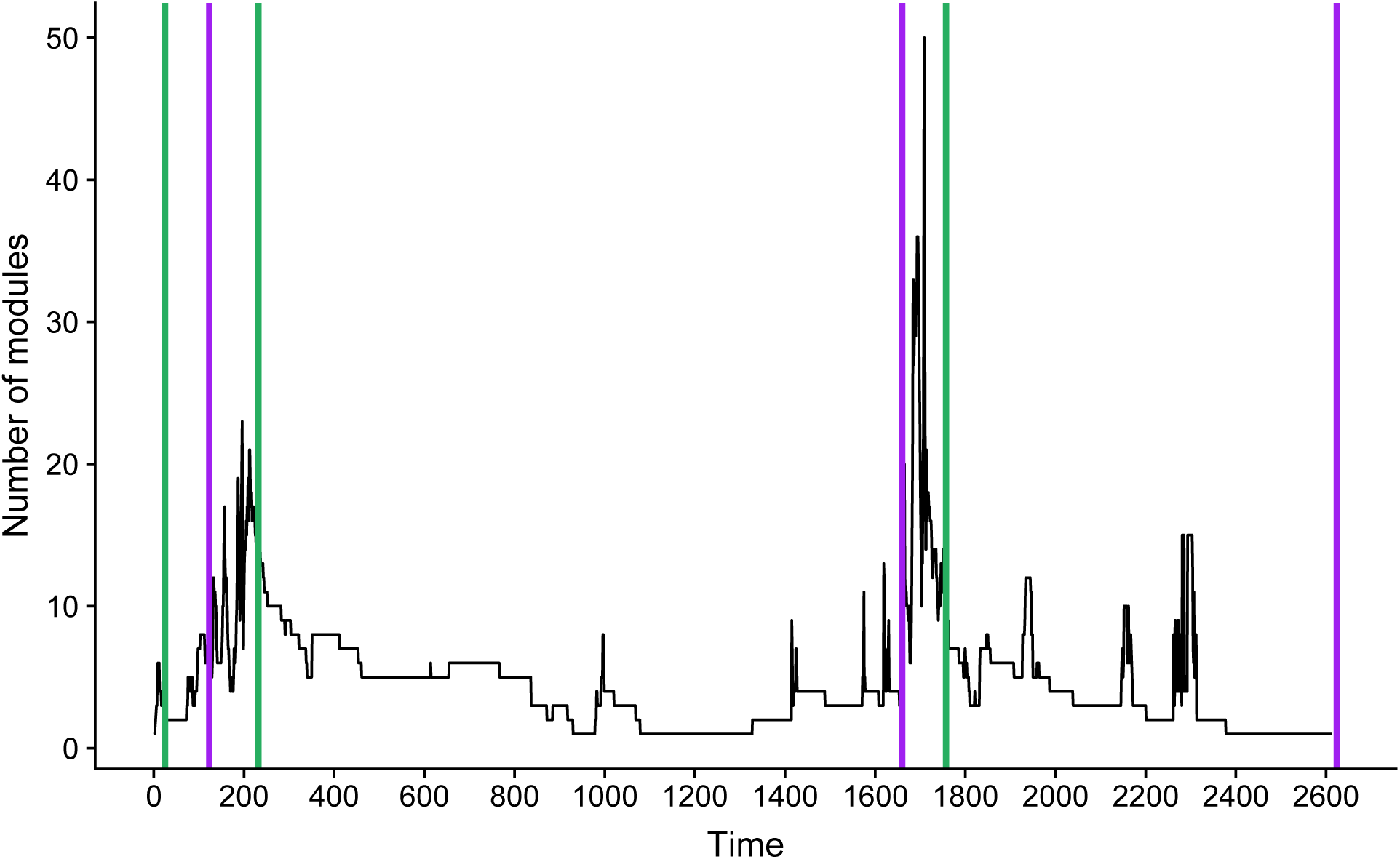
Modules in the infection network. A time series of the number of modules in the infection network for a single simulation. Modularity enables diversification since it allows the temporary coexistence of different groups of viruses.

**Fig. S5.**
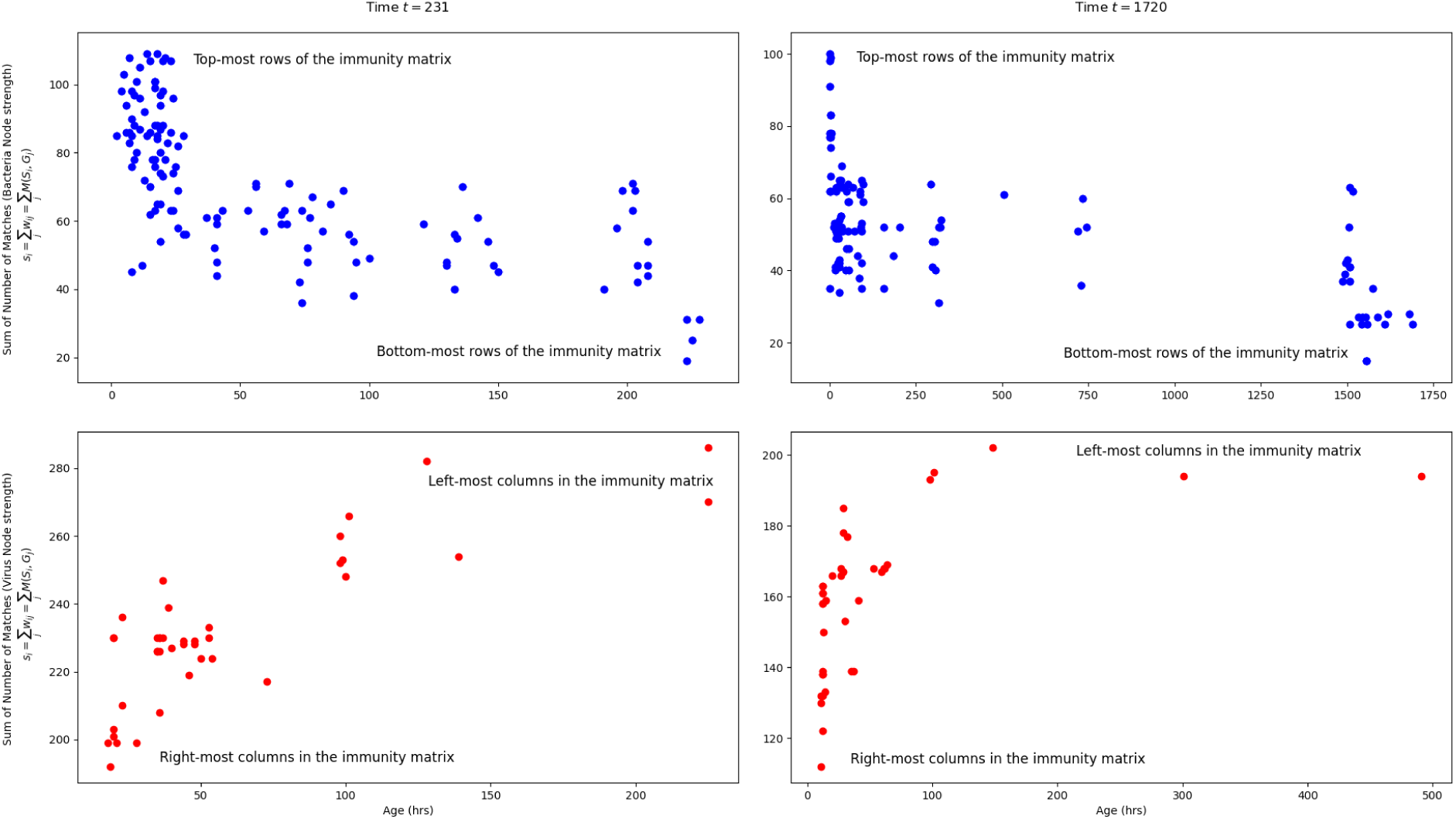
Host and virus rankings in the weighted nested immunity matrix as a function of age. The plots show node strength (sum of the corresponding number of matches in the immunity network) of hosts (top) and viruses (bottom) against their age measured from the time of their birth, for two selected times before the start of an HCR (left, t=231 and right, t=1720). Each data point represents a host (top) or virus (bottom) strain. On the different plots, selected groups of youngest and oldest strains are indicated. The oldest host strains occupy the lower rows (low node strength), and their rankings tend to decrease with age. This is because descendants of a given host inherit all of its spacers and add a new one, which always results in an increase in total matches (host node strength). Because they have acquired additional protection, they can grow in abundance and through resulting enhanced encounters and infections, the failure of existing spacers can add redundancy (more than one spacer to the same virus), further contributing to their ranking. The oldest viruses occupy the leftmost columns, with the highest column sums of matches to hosts, since longer lifetimes provide the opportunity for many encounters, and therefore for both the failure of existing spacers (which adds redundancy to a given entry) and the addition of new spacers (which distributes immunity throughout entries). A successful offspring will have mutated a protospacer that confers escape from a given match of the parent; thus, successful descendants exhibit one less match and are placed to the right. It is worth noting that there is considerable variation around the general trends with age, reflecting the complex interplay of the stochastic acquisition of spacers and protospacers with the abundance dynamics which affect both encounter and mutation rates.

**Fig. S6.**
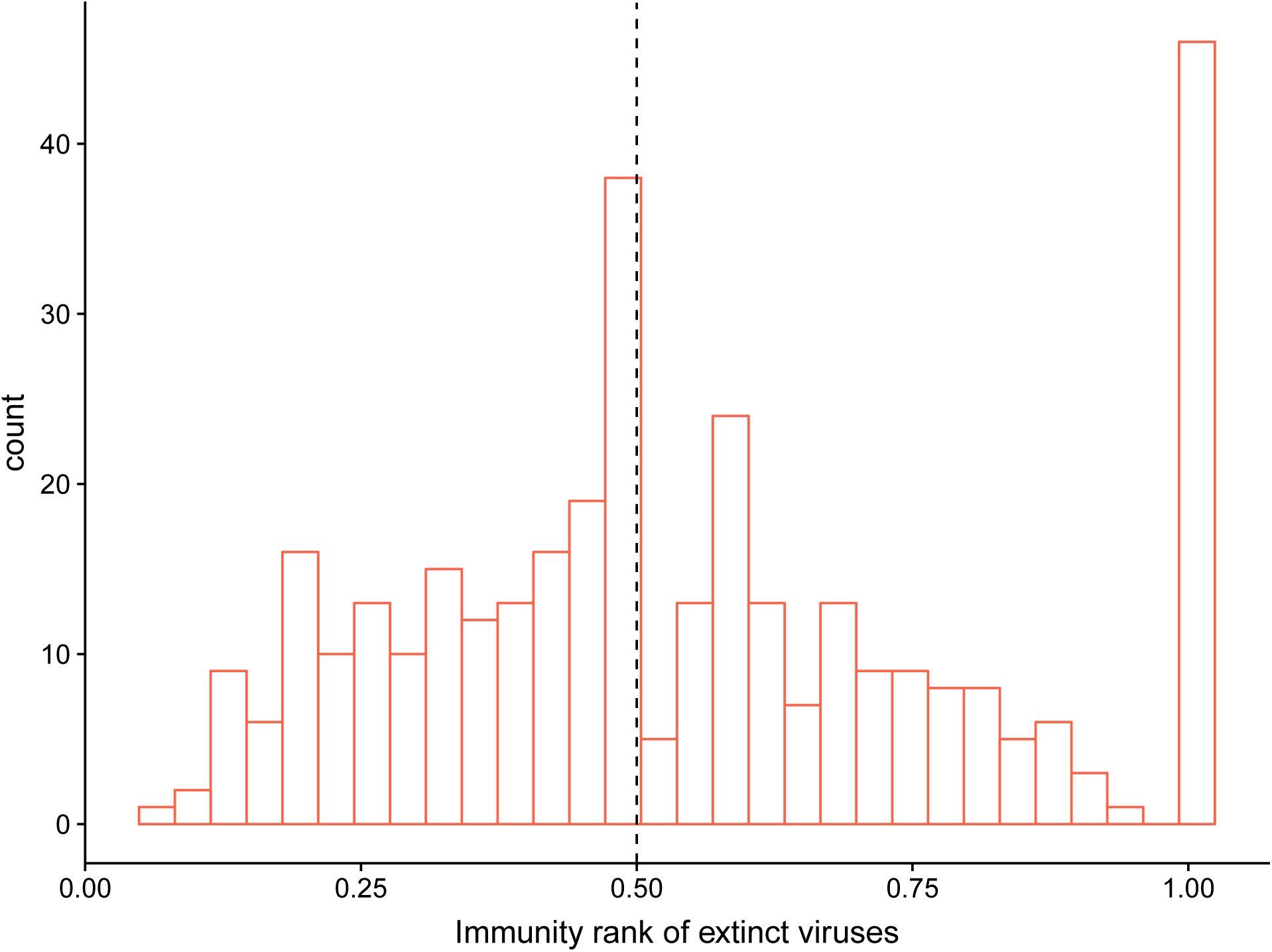
Order of extinctions. We tested for “orderly” extinctions in which extinction preferentially happens from the viruses to which hosts have most immunity to those that can infect many hosts. For any virus that went extinct we calculated an ‘immunity rank’. Specifically, for a given time step, we calculated the strength of all *n* virus nodes in the immunity network, *s* = (*s*_1_, *s*_2_, …, *s*_*j*_, …, *s*_*n*_), where *s*_*j*_ is the node strength of virus *j* (i.e., the sum of the columns in Fig 2B). Viruses with higher values of *s*_*j*_ are those that are more to the left in Fig 2B, and to which hosts have high immunity. We removed duplicate values in *s* (to avoid ties) and ordered it in ascending order to obtain *s*′. We then calculated the relative position of *s*_*j*_ in *s*′. A rank of 1 means that the virus that went extinct was highly ranked (e.g., position 5 out of 5 values will render a rank of 1). 50% of viruses (median indicated by a vertical dashed line) had an extinction rank of 0.5.

**Fig. S7.**
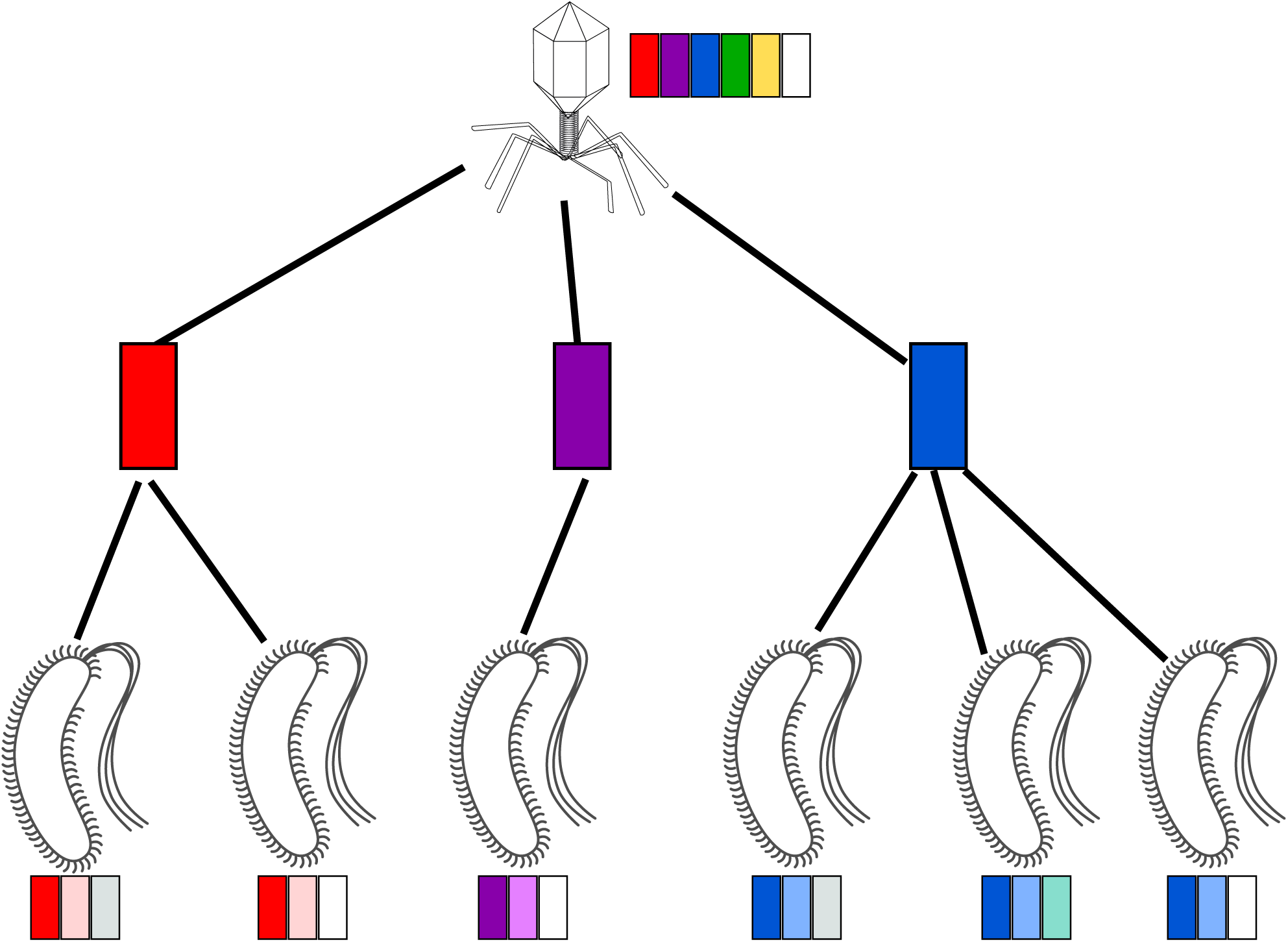
Viral escape via 1-matches. A tripartite virus-protospacer-host network depicting escape routes for a single virus. Each host is connected to a single protospacer (colored boxes). The spacer composition of strains is shown. Escape occurs through matching colors.

**Fig. S8.**
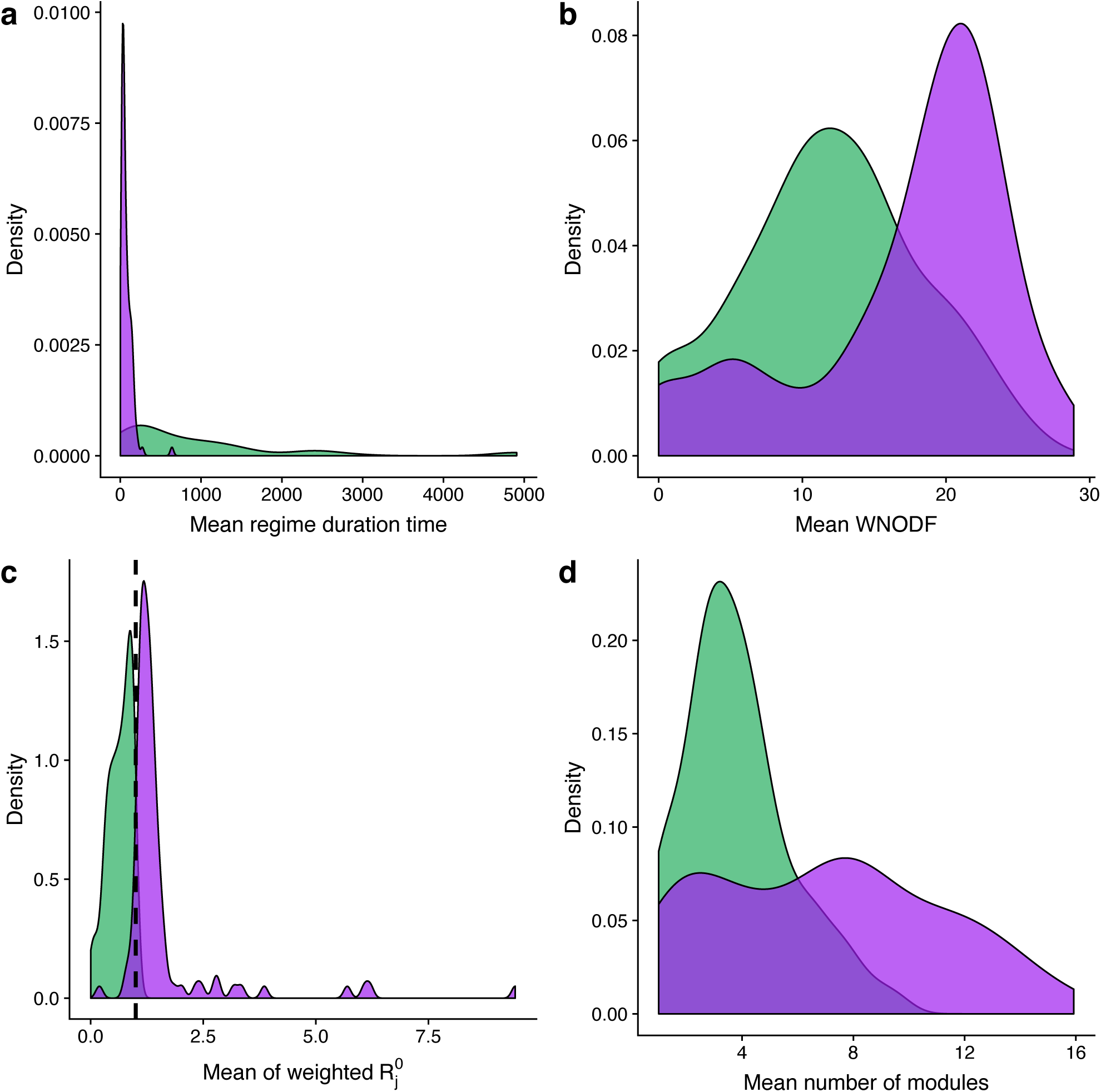
Results for multiple simulations comparing VDRs to HCRs. Results are summarized using distributions of the main characteristics of the regimes and their corresponding network structures. **a** Regimes length: VDRs are shorter than HCRs. **b** Nestedness is higher during VDRs than HCRs. **c** The basic reproductive number is higher during HCRs, indicating that virus reproduction is higher. During HCRs the basic reproductive number is generally small than one (dashed vertical line). **d** The infection network has more modules during VDRs due to virus diversification. In all plots the average was first taken for each regime type (VDR or HCR) within each simulation. Then, these averages were plotted as a distribution of 100 simulations. VDRs and HCRs are in purple and green, respectively.

**Fig. S9.**
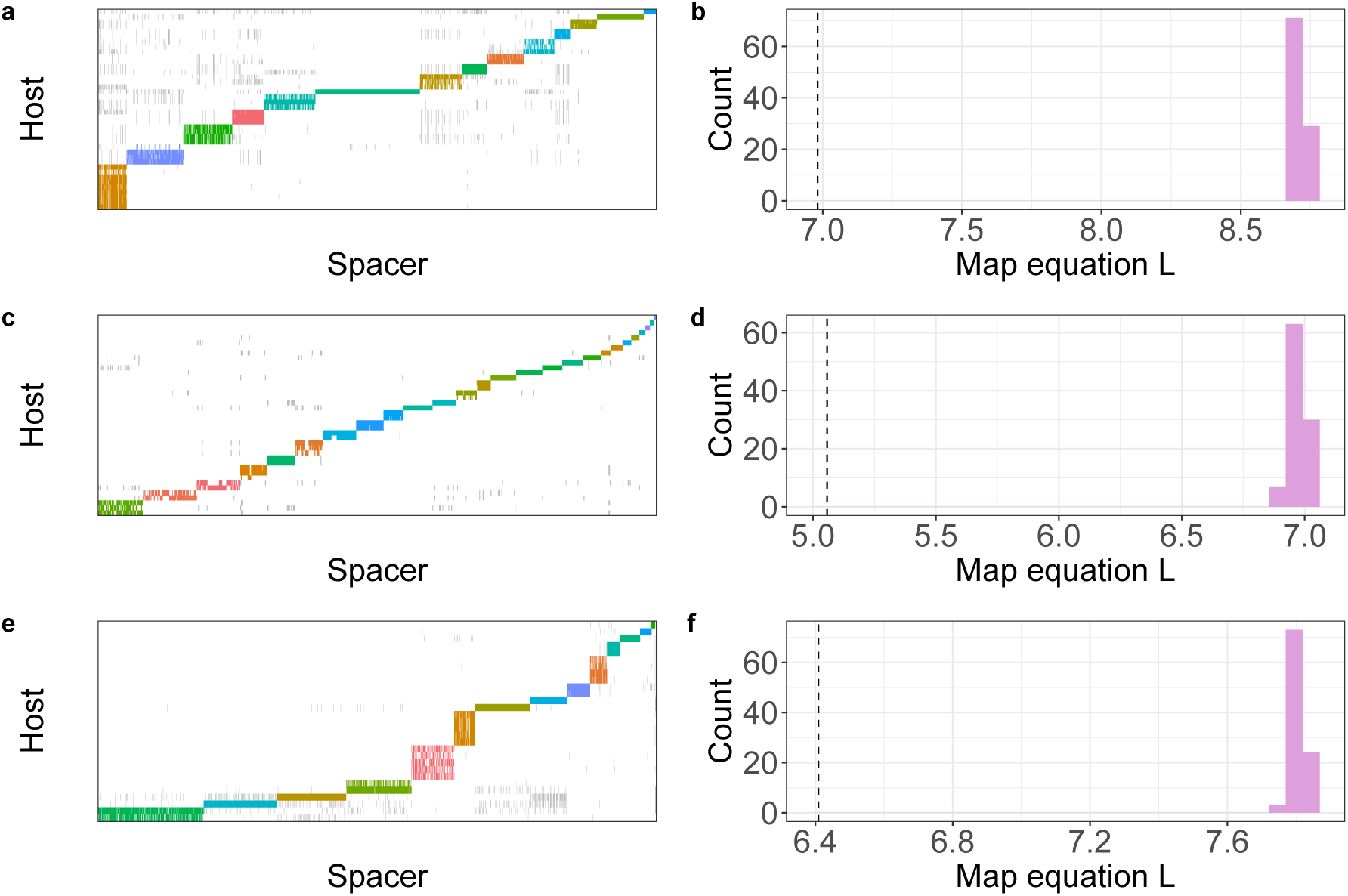
Modularity of empirical host-spacer networks. Each row represents a different data set: *Sulfolobus islandicus* hosts from Yellowstone (Top). *Pseudomonas aeruginosa* hosts from Copenhagen (middle). *S. islandicus* hosts from the Mutnovsky Volcano in Russia, 2010 (bottom). Panels **a, c** and **e** are host-spacer networks in which interactions within host-spacer modules are colored. Panels **b, d** and **f** are distributions of the map equation (*L*) obtained from networks shuffled by randomly distributing interactions. Value of the observed *L* is depicted with a vertical dashed line.

**Fig. S10.**
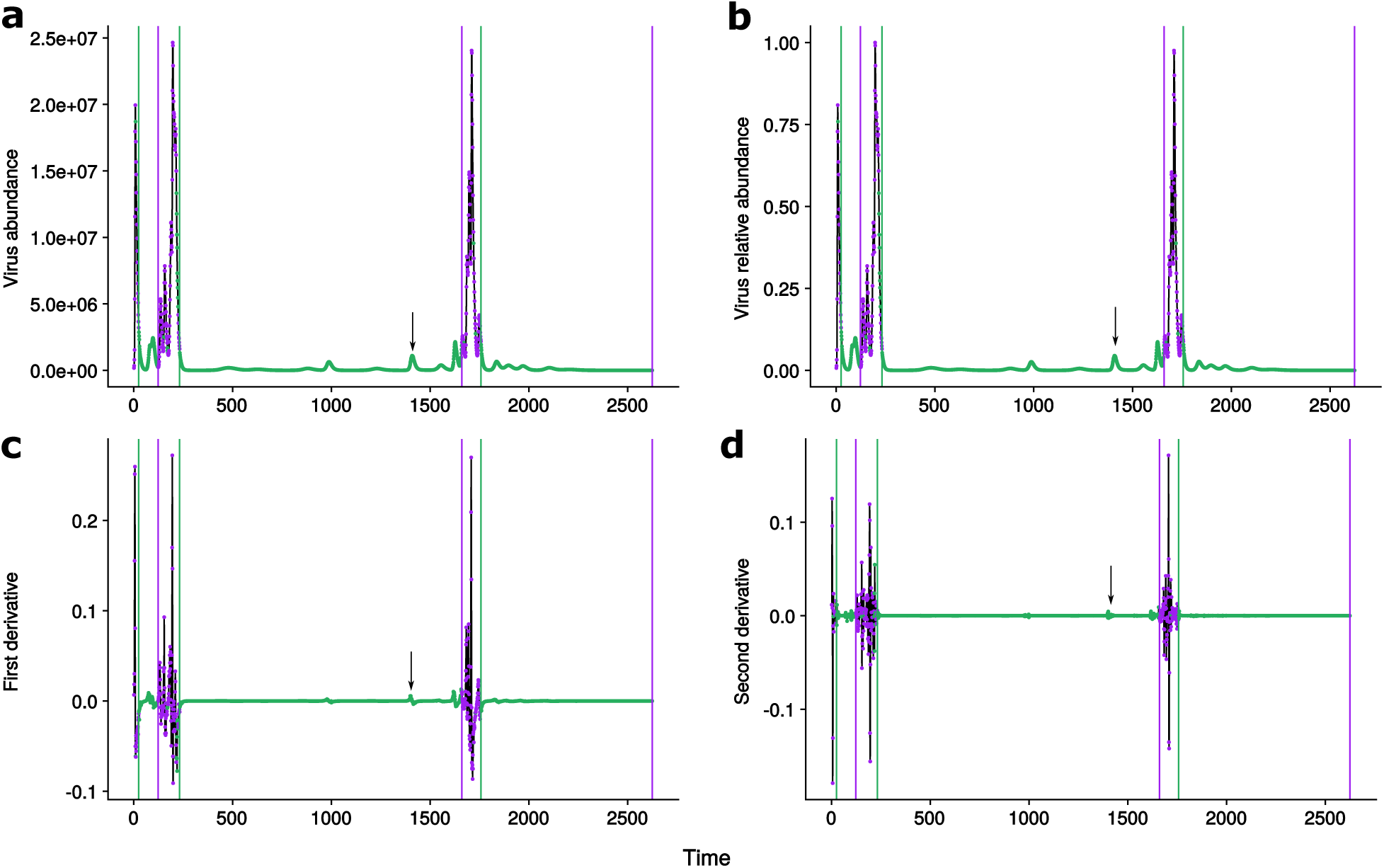
Regime definition. Each point in the virus abundance time series in panel **a** is first converted to relative abundance (panel **b**) and then classified into a HCR (green) or VDR (purple). This classification is based on the second derivative (panel **d**) (for comparison we also show the first derivative in panel **c**). Momentary virus growth periods (marked with an arrow) are not classified as VDR. The final classification is shown using vertical lines. HCRs start with a purple line and VDRs with a green line.

## Supplementary Methods

### A Implementation of dynamical model

We used a Gillespie algorithm to introduce stochasticity in spacer acquisition and protospacer mutation. The Gillespie algorithm was originally conceived for chemical reactions^1^ and is now widely used in epidemiology and ecology (e.g.^2–5^). The basic algorithm consists of three steps: (1) determination of the random time to the next reaction (also termed ‘event’); (2) random selection of a specific event from a set of events using weighted sampling (events with higher rates are more likely to be chosen); and (3) updating the number of chemical molecules (in our case, host or virus strains) involved in the selected event. Mathematically, consider *N* stochastic events with corresponding rates *a*_1_, *a*_2_, …, *a*_*N*_. The random time to an event is given by 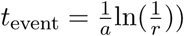, where 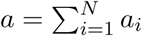 and *r* is a random number uniformly distributed in the range [0, 1). A specific event is selected with probabilities proportional to its rate, and the corresponding state numbers concerning this event are updated.

In the CRISPR coevolutionary model, we consider two stochastic events: spacer acquisition and protospacer mutation. To apply the Gillespie algorithm, we first calculate the spacer acquisition rates for all host strains and the mutation rates for all protospacers in all virus strains. We use these rates to then determine the time to the next stochastic event, to select a virus or host strain for the given change, and to add the newly generated host (in case of spacer acquisition) or virus (in case of a mutation event) strain to the system.

Mathematically, the spacer acquisition rate for host strain *i* is *a*_*i*_ = *qϕ* ∑_*j*_ *N*_*i*_*V*_*j*_ and the protospacer mutation rate for protospacer *y* of virus strain *j* is *µ*_*jy*_ = *µβϕ*(1 − *q*) ∑_*i*_ *N*_*i*_*V*_*j*_(1 − *M*_*ij*_) + *µβϕp* ∑_*i*_ *N*_*i*_*V*_*j*_*M*_*ij*_. The random time to the next stochastic event is 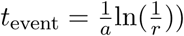 where *a* = ∑_*i*_ *a*_*i*_ + ∑_*j*_ ∑_*y*_ *µ*_*jy*_ and *r* is a random number uniformly distributed in the range [0, 1). With probability 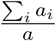, this process randomly selects a specific host to acquire a spacer from a random virus strain (with probabilities proportional to their spacer acquistion rates). Otherwise, this process randomly selects a specific protospacer in a specific virus strain to be mutated (with probabilities proportional to their mutation rates), generating a new virus strain.

This simulation scheme adopts the deterministic treatment of population dynamics in between stochastic events and follows the protocol proposed in^4^, where more details can be found. Population dynamics could also be implemented stochastically as in^6^. This would add demographic stochasticity to population dynamics, and in particular to the processes of extinction and mutant emergence. A comparison of a full stochastic implementation to the hybrid one used here is underway.

### B *R*_0_ expression

We derive here the general expression for the basic reproductive number (*R*_0_) of the system. The growth rate of a virus strain *j* is given by:

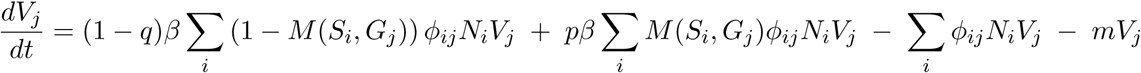

where *M* (*S*_*i*_, *G*_*j*_), or more simply *M*_*ij*_, is the number of matches between the set of spacers of bacteria *i* and the set of protospacers of virus *j*.

The condition for the spread of this virus strain 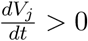 becomes

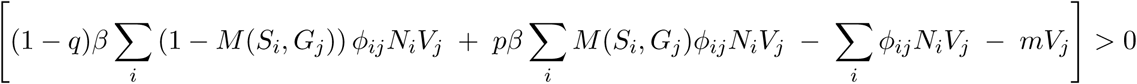

or

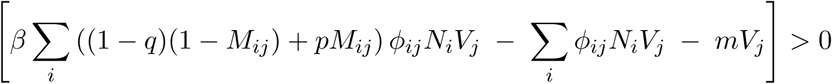

By assuming that *ϕ*_*ij*_ = *ϕ* is constant and the same for all strains,

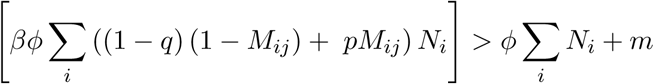

Then,

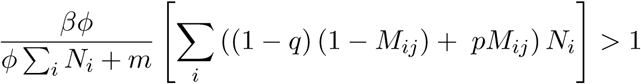

With the total abundance of bacteria *N*_*T*_ = ∑_*i*_*N*_*i*_,

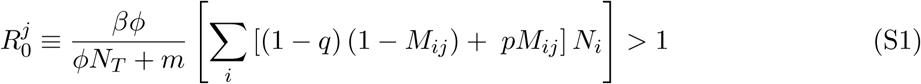

Thus,

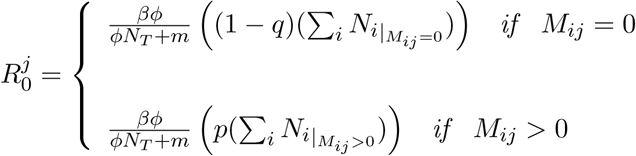

We can call the first term of equation S1 the effective *R*_0_, or 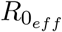, resulting from feasible infections of bacteria that do not have immunity to the given virus. The second term can be neglected since the probability of CRISPR failure *p* ≪ 1.

### C Potential *R*_0_ resulting from an escape mutation

We define the potential reproductive number of a virus as the number of offspring it would produce from a single escape mutation. For this, we need to consider the contribution to this number from infections of bacteria that are protected by a single match, 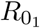. We let *g*_*j*_ denote the number of protospacers per strain and write

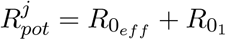

Then,

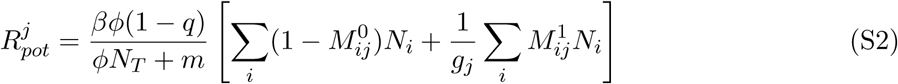

where 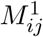 is 1 only when there is a single match between the pair of virus *j* and bacteria *i*, and 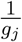 is the probability that the mutation falls on a particular protospacer of virus *j*.

### D Lotka-Volterra mean-field dynamics

The simplest model whose dynamics can be compared to those of the full model corresponds to the mean-field differential equations describing the bacteria-virus interactions when the structure of who is protected from whom is randomized. That is, in this model, the emergent structure of the protection matrix is randomized, and the immunity matrix is reduced to a fixed density of edges immunity. Starting from the full model from Childs et al.,

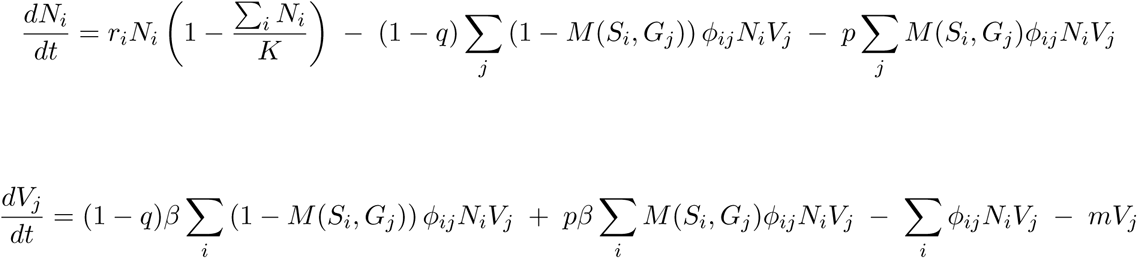

and simplifying some notation, we have

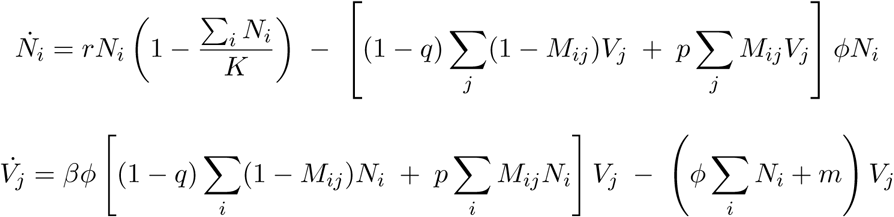

We assume that the entries of the immunity matrix *M*_*ij*_ (defined by the matches between the set of spacers (*S*_*i*_) of the host *i* and the set of protospacers (*G*_*j*_) of the virus *j*) take a constant value in time, *M* ≡ ⟨*M*_*ij*_⟩. We further rewrite the system for the total abundance of virus *V* and bacteria *N* as the following general predator-prey system:

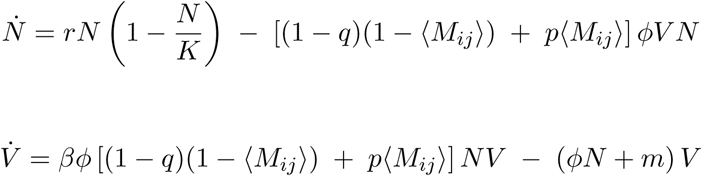

With 𝕄 ≡ [(1 − *q*)(1 − ⟨*M*_*ij*_⟩) + *p*⟨*M*_*ij*_⟩], we can rewrite the system as:

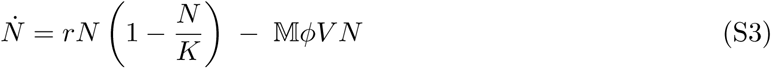

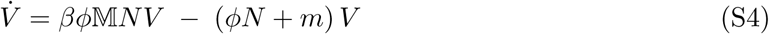

#### D.1 Equilibrium points

The above two-dimensional system (S3 and S4) has 3 equilibrium points *C*_*i*_ = (*N* ^∗^, *V*^∗^):

- The trivial point *C*_1_ = (0, 0).
- The point where hosts have reached their carrying capacity and the viruses have gone extinct, *C*_2_ = (*K*, 0).
- The positive coexistence point defined by the intersection of the nullclines of the system

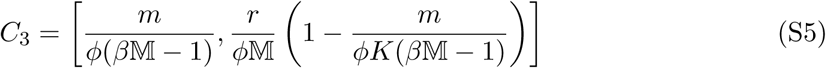

These nullclines are both a function of 𝕄 and are given by:

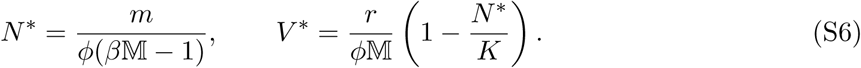

The local (asymptotic) stability of these equilibria can be determined based on the Jacobian matrix of the system :

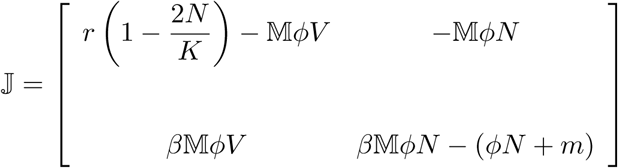

Together the expressions for the nullclines and for the Jacobian show that the location of the equilibria in phase space and their local stability depend on 𝕄 and therefore, on *M* = ⟨*M*_*ij*_⟩. The change in the equilibria as a function of *M* is illustrated in Supp Methods Fig. 1. These dynamics can be compared to those of the full system for the aggregated abundance of all viruses and hosts (Supp Methods Fig. 2).

**Supp Methods Fig. 1.**
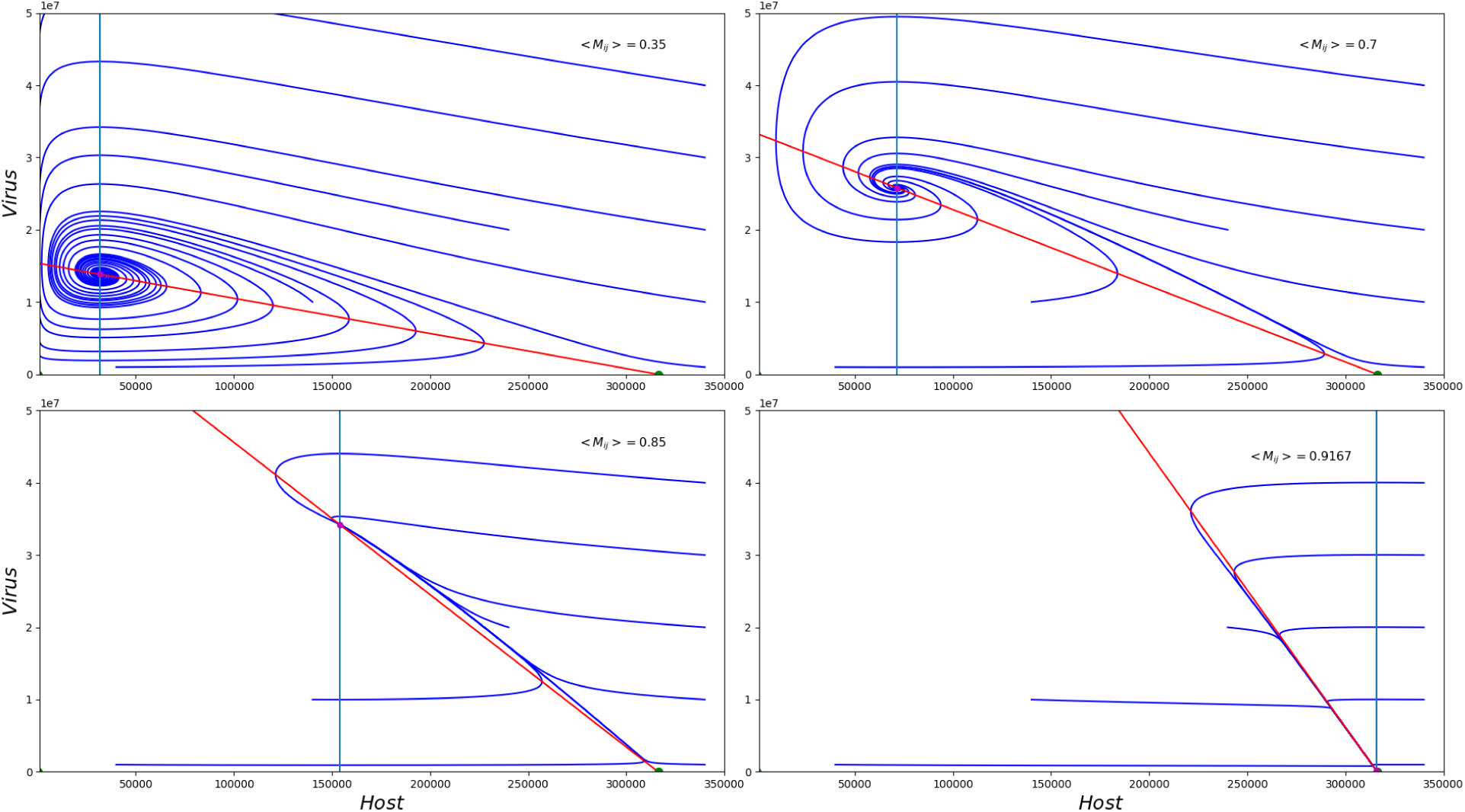
Phase portraits of the mean-field ODE version of the system for increasing values of the average fraction of matches *M*. Top panels from left to right: *M* = 0.35 and *M* = 0.7; bottom panels (also from left to right): *M* = 0.85 and *M* = 0.9167. As *M* grows, the fixed point *C*_3_ transitions from a spiral sink to a degenerate nodal sink as it approaches *C*_2_ and finally collapses onto it.

### E Neutral model without explicit immunity (Randomization of the immunity matrix)

By construction, the above mean-field simplification of the system does not explicitly include the diversity of hosts and viruses. To determine the importance of both specific immune memory and the structure associated with it, we consider here a neutral model in which the ability of the system to generate diversity is not affected (that is, both the mutation of viruses and the acquisition of spacers by the bacteria are still included) but the specificity of immune memory is removed. We specifically randomize every Δ*t* the matches *M* (*S*_*i*_, *G*_*j*_) between the protospacers set of viruses (*G*_*j*_) and the spacers set of hosts (*S*_*i*_). For every virus-host strain pair, we randomize their match by choosing a random number *r* and comparing it with the *average density* of the immunity network, *ρ*. We define the density of a an infection network at time *t* as *C* = *E/vn*, where *E* is the total number of infection interactions and *v* and *n* represent the number of virus and host strains, respectively. The average density is then the average of *C* across all time points. Then, we establish a match for this pair if *r* < *ρ*. This preserves the average density of matches but randomizes their identity.

**Supp Methods Fig. 2.**
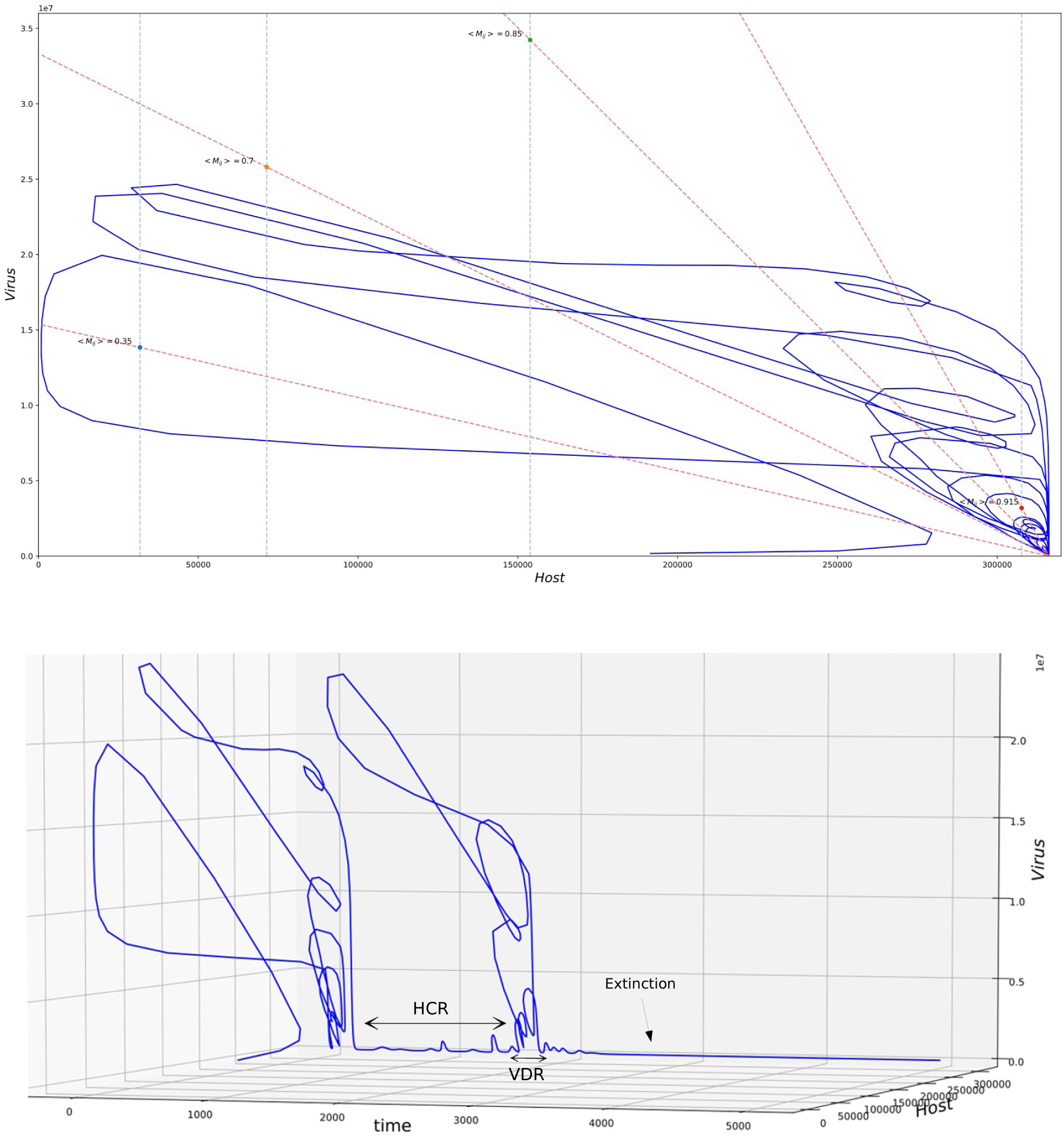
Phase portrait of the full system for total abundances of hosts and viruses. The dynamics of the full system differ from those of the two-dimensional mean-field ODE as expected from the transitions from coexistence (during the VDR) and the host dominance at carrying capacity (during the HCR) (top panel). During the VDR, the system’s trajectory spirals around a moving coexistence point, closest to the fixed points *C*_3_ for increasing *M* (color points). During the HCR, the system is close to the degenerate sink, collapsing onto *C*_2_ after virus extinction. These different parts of the dynamics become more evident when including time explicitly in the phase portrait (bottom panel).

Under this null model, the system shows neither the regime of virus diversification (VDR) nor that of host dominance (HCR) (results not shown). This comparison to the full model established that the structure of specific immunity generated by the CRISPR system is necessary to create the observed dynamics of the full model.

### F Detailed methods for empirical data collection

Our data consists of three data sets:

#### Yellowstone

This dataset was collected in hot springs in Nymph Lake at Yellowstone National Park and consists of a population of *Sulfolobus islandicus* and its contemporary lytic (Sulfolobus islandicus rod-shaped viruses: SIRVs).

#### Russia

This dataset consists of a set of *S. islandicus* strains isolated from Kamchatka, Russia (in 2000) and sympatric chronic viruses (Sulfulobus spindle-shaped viruses: SSVs)^7^.

#### Pseudomonas

This dataset consists of longitudinal sampling of human-adapted *Pseudomonas aeruginosa* isolates from sputum samples of Cystic Fibrosis patients collected at a hospital in Copenhagen, Denmark and a global set of temperate and lytic *P. aeruginosa* viruses extracted from NCBI to substitute for a lack of sequenced contemporary viruses^8–10^. Viruses were grouped based on nucleotide similarity into families known as clusters, and these clusters were assigned a number identifiers which have been described in a previous study (4). In this study we only used viruses from cluster 3 to avoid a false positive result in which we obtain a nested structure due to immune patterns that depend on the cluster (phylogeny).

Illumina sequenced reads from samples were quality filtered using prinseq with the following arguments: -derep 1245 -lc_method entropy -lc_threshold 50 -trim_qual_right 30 -trim_qual_left 30 -trim_qual_type min -trim_qual_rule lt -trim_qual_window 5 -trim_qual_step 1 -trim_tail_left 5 -trim_tail_right 5 -min_len 66 -min_qual_mean 30 -ns_max_p 1 -verbose. Spacers were extracted from quality filtered sequencing reads using an in-house bioinformatic pipeline (in preparation, code available upon request). These scripts utilize known repeats from *S. islandicus* and *P. aeruginosa* and BLASTn to extract spacers from reads located between any repeats. BLASTn cutoffs for repeats against reads were based on an e-value of with the `-task blastn-short argument`. After spacer extraction, spacers are grouped based on a hamming distance cutoff by comparing spacers as strings of nucleotides and using a sliding window across each string of basepairs (in preparation, code available upon request). We define unique spacers that have 100% nucleotide identity to one another, using a hamming distance of 0 between nucleotide sequences. Unmatched overhanging base pairs between spacers were considered mismatches since these are likely independently acquired spacers from sequential protospacers with different PAM sequences. Unique spacers were mapped to strains in order to determine the spacer set per strain. *Sulfolobus islandicus* isolates in Nymph Lake contained on average 256 spacers per strain ranging from 62 to 520 spacers. *Sulfolobus islandicus* isolates from Kamchatka contained on average 181 spacers per strain ranging from 20 to 795 spacers. *P. aeruginosa* isolates contained 34 spacers on average with a range of 4 to 64 spacers. *P. aeruginosa*, Nymph Lake *S. islandicus*, and Mutnovsky *S. islandicus* isolates contained 40, 40 and 50 total alleles respectively.

Spacer matches to protospacers were initially found using BLASTn with a `-task blastn-short` argument, with an e-value minimum of 0.01. The *P. aeruginosa* database contained 6,231,702 total bp, with 98 viral genomes ranging from 3,588 bp to 309,208 bp. The SSV BLASTn database contained 34 genomes containing 514,147 total bases with the longest sequence being 18,548 bp and the shortest being 11,323^10,11^. The SIRV BLASTn database was composed of 10 genomes containing 347,896 total bases with the longest sequence being 32,308 bp and the shortest being 32,308. Protospacer BLAST matches were extended to 3 base pairs longer than the length of the spacer and retrieved with blastdbcmd tool from the blast+ package to retrieve the PAM sequences^12,13^. Gaps found between alignments were added to these extended protospacer matches. Gaps, or insertion/deletion events, were considered as mismatch when comparing along the entire length of the aligned protospacer and spacer.

We analyzed our data with two criteria. The first criterion was a perfect match with zero mismatches between the entire length of the aligned spacer and protospacer (100% alignment) and with correct PAM sequences in the correct orientation based on the 3bp extensions. The range in number of protospacers found in *S. islandicus* viruses was 39-41 in SIRVs with and 11-47 in SSVs at the most stringent criterion. The range of protospacers found in viruses was 0-32 with *P. aeruginosa* viral database. In our second criterion, we allowed 4 mismatches and do not require a specific PAM match. The number of protospacer matches to viruses ranged from 138 to 167 with SIRVs and 6-24 in SSV for 4mm criterion. Protospacer matches ranged from 0-76 with *P. aeruginosa* viruses. Our results were not qualitatively affected by the choice of criterion and we present results for the second one.

